# Ubiquitination of G3BP1 mediates stress granule disassembly in a stress-specific manner

**DOI:** 10.1101/2021.04.22.440930

**Authors:** Youngdae Gwon, Brian A. Maxwell, Regina-Maria. Kolaitis, Peipei Zhang, Hong Joo Kim, J. Paul Taylor

## Abstract

Stress granules are dynamic, reversible condensates composed of RNA and protein that assemble in response to a variety of stressors and are normally disassembled after stress is removed. Whereas the composition of stress granules and the mechanisms underlying their assembly have been extensively studied, far less is known about the mechanisms that govern disassembly. Impaired disassembly has been implicated in some diseases. Here we report that stress granule disassembly is context-dependent and, in the setting of heat shock, requires ubiquitination of G3BP1, the central protein within the stress granule RNA-protein network. Ubiquitinated G3BP1 interacts with the ER-resident protein FAF2, which engages the ubiquitin-dependent segregase p97/VCP. Targeting G3BP1 enables the stress granule-specific interaction network to fall below the percolation threshold for phase separation, which causes disassembly.

**One Sentence Summary:** Ubiquitination of G3BP1 mediates FAF2- and p97/VCP-dependent disassembly of heat-induced stress granules

## Main Text

Biomolecular condensation is a vital strategy of cellular organization that regulates a wide variety of biological functions. Ribonucleoprotein (RNP) granules are an important and highly conserved class of biomolecular condensates that govern many aspects of RNA metabolism, including RNA biogenesis, modification, utilization, and degradation (*1, 2*). One prominent type of RNP granule is the stress granule, a dynamic and reversible cytoplasmic assembly formed in eukaryotic cells in response to a wide variety of stressors. Formation of stress granules typically follows upon inhibited translation initiation and polysome disassembly, which gives rise to a rapid increase in cytoplasmic concentration of uncoated mRNA. This rise in cytoplasmic mRNA triggers multicomponent liquid-liquid phase separation (LLPS) with RNA-binding proteins, creating a condensed liquid compartment that remains in dynamic equilibrium with, yet distinct from, the surrounding cytosol (*3–5*). Impaired dynamics of RNP granules such as stress granules are implicated as an important contributor to certain pathological conditions, including neurodegenerative diseases (*6*).

The mechanisms underlying assembly of biomolecular condensates have been a subject of intense interest, and recent investigations of stress granule assembly have yielded significant insight into how multicomponent networks self-organize to form a compartment that is distinct from its surroundings. At the level of individual molecules, stress granules are composed of RNA and protein that interact via protein protein-protein, protein-RNA, and RNA-RNA interactions, each of which may be weak or transient. However, once the sum of these interactions breaches a particular threshold, known as the percolation threshold, the individual molecules form a system-spanning network that separates itself from its milieu – in other words, LLPS ensues (*3–5*), creating two distinct phases: a dense phase enriched in macromolecules (i.e., the stress granule) that coexists with a dilute phase that is deficient in macromolecules (i.e., the cytoplasm). The concept of a percolation threshold is generalizable to other biomolecular condensates, each of which encodes a system-specific threshold for LLPS.

In stress granules, there are ~36 proteins that, together with RNA, provide the majority of the interactions that set the percolation threshold for RNA-dependent LLPS (*3*). Whereas each constituent node of this network contributes toward the sum of interactions required to reach the percolation threshold, a small subset of “scaffolding” constituents dominate in establishing this threshold (*3, 7*). The most important proteins in the stress granule interaction network are G3BP1 and G3BP2, and these paralogous proteins provide the largest contribution to establishing the percolation threshold for stress granule assembly (*3–5*). Indeed, the importance of G3BP1/2 proteins is such that their elimination precludes stress granule assembly in some contexts (*3, 8*) and their enforced activation is sufficient to initiate stress granule assembly even in the absence of stress (*3, 9*). Higher order regulation over stress granule assembly may also be afforded by post-translational modifications of stress granule proteins that impact the interaction network, but the relative importance of individual types of modifications and their specific targets remains to be determined (*10*).

In healthy cells, stress granules are transient, dynamic structures whose presence in the cell generally corresponds to the duration of time during which a particular stress is applied. However, exactly how stress granules are removed from cells after stress is relieved remains an unresolved question. Indeed, untangling the current literature regarding stress granule elimination first requires careful consideration of the duration and the specific type of stress applied. The duration of stress largely dictates whether stress granules are eliminated via *clearance* versus *disassembly*. During prolonged stress, or in the setting of disease mutations, stress granules become gradually less dynamic and are eventually *cleared* through an autophagy-dependent degradative process that requires VCP and autophagy machinery such as ATG7 (*6, 11, 12*). Importantly, autophagy-dependent clearance of stress granules results in degradation and permanent loss of constituent proteins and RNAs. In contrast, if cellular stress is removed relatively quickly, while stress granules remain dynamic, they are *disassembled* by decondensation – a reversal of LLPS – that occurs when sum of protein-protein, protein-RNA, and RNA-RNA interactions falls below the percolation threshold. This decrease in the sum total of interactions within the stress granule network could occur through several mechanisms, including a decrease in the concentration of individual constituents or altered post-translational modifications of constituent proteins that weakens the interaction network. In contrast to stress granule clearance, disassembly is a non-degradative process wherein the individual constituents disassociate from one another and are recycled, including liberation of the mRNAs to re-join the translating pool (*13*).

Even within the specific route of disassembly, context matters: as shown in our companion paper (Maxwell et al., co-submitted), stress granules formed by heat stress are disassembled via ubiquitin-dependent mechanisms, whereas those formed in response to arsenite are not. Thus, the key nodes and cellular processes engaged in disassembly of stress granules are specific to each type of stress. This finding may explain why previous studies have reported that ubiquitination is dispensable for arsenite-induced stress granule dynamics (*14*) whereas others have reported that the ubiquitin-binding adaptor protein ZFAND1 is required (*15*). Another layer of complexity is demonstrated by studies of the role of VCP, a ubiquitin-dependent segregase that uses adaptor proteins to engage polyubiquitinated clients and aid their energy-dependent extraction from higher order complexes (*16*). VCP has long been implicated in stress granule clearance via autophagy-dependent mechanisms (*12*), but has also been identified in ZFAND1-mediated stress granule removal (*15*) and more recently was demonstrated to function in autophagy-independent stress granule disassembly (*17*).

Here we address these earlier apparent inconsistencies by revealing in detail the molecular mechanism of ubiquitin-dependent stress granule disassembly in the context of heat shock. Building from the observation from Maxwell et al. (co-submitted) that the central stress granule scaffolding protein G3BP1 is ubiquitinated in response specifically to heat stress, we demonstrate that context-dependent ubiquitination of G3BP1 mediates interaction with the ubiquitin-binding protein FAF2, which serves as an adaptor protein that engages VCP for ATP-dependent extraction of G3BP1. Consistent with its central role within the stress granule network, extraction of G3BP1 drives decondensation and subsequent disassembly of stress granules. Notably, FAF2 is an ER-resident protein, demonstrating that stress granules that arise in response to heat stress are disassembled at the ER, consistent with other heat shock-dependent stress responses such as the unfolded protein response (UPR) and ER-associated degradation (ERAD). This finding also resolves unanswered mechanistic questions raised by the recent observation that stress granules associate with the ER prior to their elimination (*18*).

### G3BP1 undergoes K63-linked ubiquitination upon heat shock

Using tandem ubiquitin binding entity (TUBE) enrichment and di-Gly enrichment assays followed by label-free LC-MS/MS, we previously identified 373 proteins that showed altered ubiquitination levels in response to heat stress, a collection of proteins we designated as the “heat shock ubiquitinome” (Maxwell et al., co-submitted). In that study, we found that the heat shock ubiquitinome is highly enriched in cellular pathways activated or perturbed by stress, and that ubiquitination is essential for resumption of cellular activities following heat shock (Maxwell et al., co-submitted). One such cellular pathway centered on the formation of stress granules and subsequent disassembly during recovery. Whereas ubiquitination was not required for heat shock-induced stress granule assembly, it was essential for their rapid disassembly following removal of stress. However, the mechanism whereby ubiquitination was required for stress granule disassembly remained undefined.

Among the stress granules proteins ubiquitinated in a context-dependent manner following heat shock was G3BP1. We previously identified G3BP1 as a central node in the protein-RNA interaction network that drives the formation of stress granules (*3*). Furthermore, pharmacologically disrupting the dimerization of G3BP1 led to rapid disassembly of stress granules, underscoring the central role played by G3BP1 in assembling and maintaining the assembly of stress granules (*3*). Thus, we hypothesized that ubiquitination of G3BP1 may have a key role in this process.

To confirm that G3BP1 is ubiquitinated in response to heat shock, we performed TUBE pulldown of ubiquitinated proteins from U2OS cells exposed to heat shock (43°C) followed by immunoblotting (**Fig. 1A**), using *G3BP1/2* double knockout (dKO) cells as controls. We detected increased levels of poly-ubiquitin-conjugated G3BP1 as early as 15 minutes after exposure to heat stress; with continuing stress, these levels peaked at approximately 60 minutes and remained detectable in the soluble fraction for at least 150 minutes (**Fig. 1B**). After stress was removed, levels of poly-ubiquitinated G3BP1 decreased, returning to baseline within 3 hours after a 60-minute heat stress (**Fig. 1C**). Interestingly, poly-ubiquitin conjugation of G3BP1 was specific to heat stress: despite an increase in total poly-ubiquitin conjugates in response to oxidative stress (sodium arsenite) or osmotic stress (sorbitol), cells exposed to these stresses did not accumulate poly-ubiquitinated G3BP1 (**Fig. 1, D and E**).

**Fig. 1.**
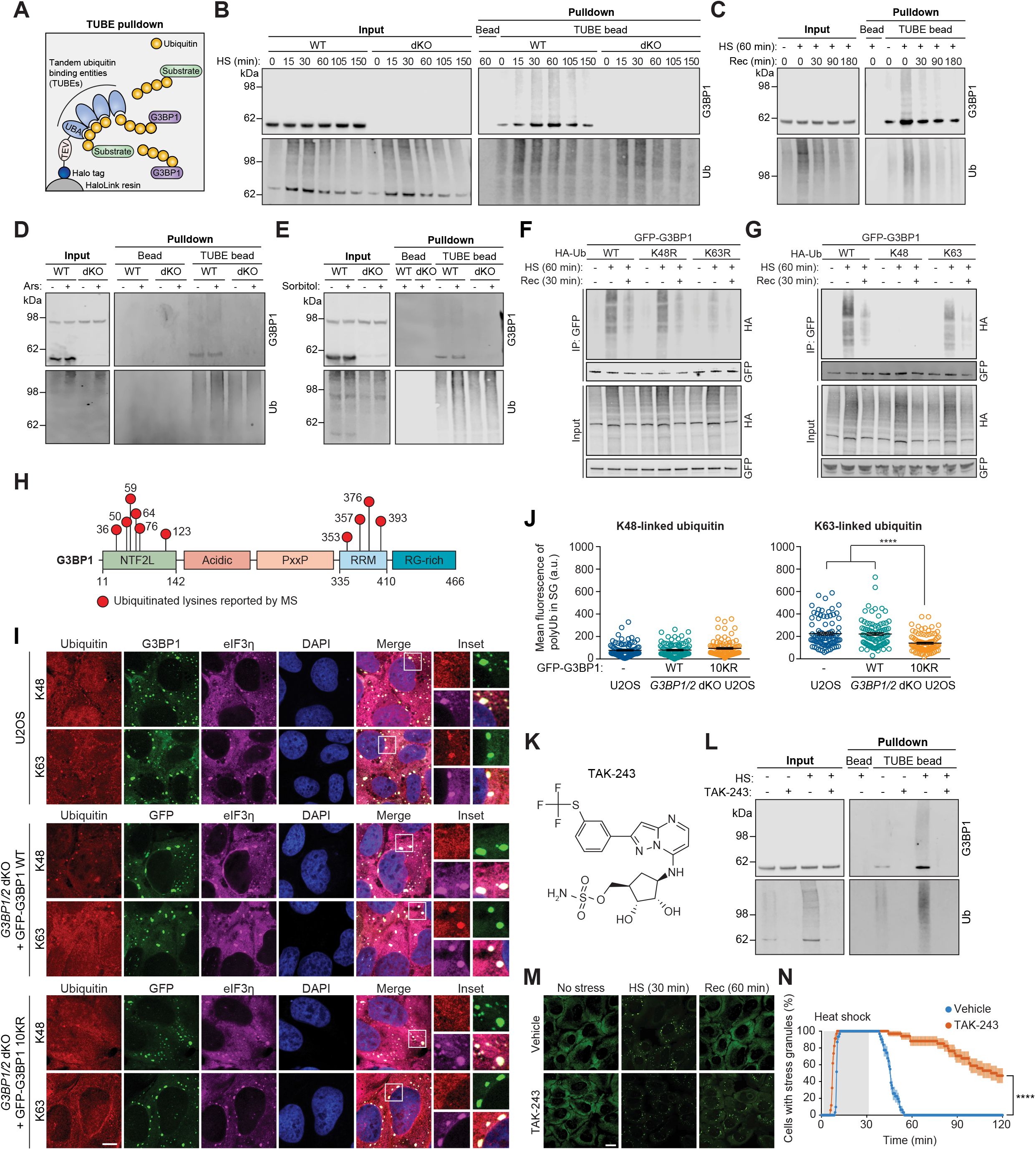
G3BP1 undergoes K63-linked ubiquitination in response to heat stress. (**A**) Schematic illustration showing TUBE capture of ubiquitinated G3BP1. (**B-C**) Western blots of TUBE-captured cell extracts showing levels of ubiquitinated G3BP1 in U2OS cells based on duration of 43°C heat shock, using U2OS *G3BP1/2* dKO cells as controls (B), and during 37°C recovery after heat shock (43°C, 1 h) (C). (**D-E**) Western blots of TUBE-captured cell extracts showing levels of ubiquitinated G3BP1 in response to oxidative stress (0.5 mM sodium arsenite, 1 h) (D) or osmotic stress (0.4 M sorbitol, 1 h) (E). (**F-G**) Western blots showing K63-linked ubiquitination of G3BP1 during heat shock (43°C, 1 h) and recovery (37°C, 30 min). HEK293T cells were transfected with GFP-G3BP1 and indicated HA-tagged ubiquitin WT or mutant constructs. Cell extracts were captured with magnetic beads conjugated with GFP antibody for IP and resulting beads were analyzed by immunoblot. (**H**) G3BP1 domain organization labeled with lysine residues on which ubiquitination has been reported. (**I-J**) Fluorescent staining of U2OS WT cells and U2OS *G3BP1/2* dKO cells stably expressing GFP-G3BP WT and ubiquitination-defective mutant10KR). Cells were exposed to heat shock (43°C, 1 h), fixed, and stained with indicated linkage-specific antibodies (I), which demonstrate decreased recruitment of K63-linked ubiquitin chains to stress granules generated by ubiquitination-defective G3BP1. Fluorescent intensities of K48-linked ubiquitin (left) and K63-linked ubiquitin (right) in eIF3η-positive stress granules are plotted in (J). Scale bar, 10 μm. Error bars indicate s.e.m. \*\*\*\**P*< 0.0001, ANOVA with Tukey’s test. (**K**) Molecular structure of TAK-243. (**L**) Western blot showing block of heat shock-induced G3BP1 ubiquitination by TAK-243. U2OS cells were treated with DMSO or TAK-243 for 60 min prior to heat shock (43°C, 1 h). Cell extracts were captured with TUBE and analyzed by immunoblot. (**M-N**) Disrupted disassembly of heat shock-induced stress granules by inhibiting ubiquitination. U2OS cells stably expressing GFP-G3BP1 were treated with DMSO or TAK-243 for 60 min prior to live cell imaging. Cells were incubated at 37°C for 2 min, 43°C for 30 min, and 37°C for 88 min, and GFP signals were monitored with 30-sec intervals. Representative still images are shown in (M). The percentage of cells with ≥ 2 stress granules is plotted in (N). Scale bar, 20 μm. Error bars indicate s.e.m. \*\*\*\**P*< 0.0001, Mantel-Cox test.

Poly-ubiquitin chains can be formed by conjugating ubiquitin to one of seven lysine residues of another ubiquitin molecule, giving rise to K6-, K11-, K27-, K29-, K33-, K48-, or K63-linked poly-ubiquitin chains (*19*). Among these, K48- and K63-linked chains are the two most abundant and functionally well-characterized linkage types (*20*). To determine which of these linkage types are present in poly-ubiquitinated G3BP1, we used ubiquitin mutants that prevent the formation of K48-linked (K48R) or K63-linked (K63R) chains or that permit K48-linked or K63-linked chains exclusively (K48 or K63 being the only available lysines). Interestingly, expression of K63R inhibited accumulation of poly-ubiquitin-conjugated G3BP1 upon heat shock, whereas expression of K48R had no impact (**Fig. 1F**). Conversely, cells expressing K63 ubiquitin (in which the only available lysine is K63) accumulated poly-ubiquitin-conjugated G3BP1, whereas cells expressing K48 ubiquitin did not (**Fig. 1G**), suggesting that heat shock promotes K63-linked poly-ubiquitination of G3BP1. Notably, the levels of K63-linked poly-ubiquitinated G3BP1 were lower than the levels of poly-ubiquitinated G3BP1 using WT ubiquitin, suggesting that other types of chains also contribute to this poly-ubiquitination.

### Heat shock-induced stress granules accumulate K63-linked poly-ubiquitin chains that are required for stress granule disassembly

Examination of public databases suggests 10 lysine residues in G3BP1 that are potentially subject to ubiquitination (*21*). Of these, 6 residues are located within the N-terminal NTF2-like (NTF2L) domain and 4 in the C-terminal RNA-recognition motif (RRM) (**Fig. 1H**). When we mutated these 10 residues (G3BP1 10KR) and expressed this construct in *G3BP1/2* dKO cells, we found no ubiquitination of G3BP1 in response to heat shock, suggesting that G3BP1 10KR functions as a ubiquitination null mutant (**fig. S1A**).

We previously observed a robust increase of poly-ubiquitin signal in stress granules induced by heat stress, but not by oxidative stress (Maxwell et al., co-submitted). To examine the contributions of K48- vs. K63-linked chains to this ubiquitin signal within stress granules, we used linkage-specific antibodies. Whereas K48- and K63-linked poly-ubiquitin signals were both uniformly distributed throughout the cell under basal conditions (**fig. S1B**), only K63-linked poly-ubiquitin signals accumulated in stress granules upon heat shock in WT U2OS cells (**Fig. 1I**). Furthermore, the levels of K63-linked poly-ubiquitin signal were significantly diminished in *G3BP1/2* dKO cells expressing GFP-G3BP1 10KR, whereas *G3BP1/2* dKO cells expressing GFP-G3BP1 WT accumulated K63-linked poly-ubiquitin signal comparable to WT U2OS cells (**Fig. 1, I and J**). These results suggest that a significant portion of heat shock-induced poly-ubiquitin signal in stress granules arises from K63-linked poly-ubiquitination of G3BP1. Notably, poly-ubiquitin conjugation of G3BP1 was eliminated by treatment with TAK-243, a potent small molecule inhibitor of UBA1, a E1 ubiquitin-activating enzyme (**Fig. 1, K and L**). Consistent with our previous observations (Maxwell et al., co-submitted), blocking ubiquitination by TAK-243 significantly delayed stress granule disassembly during recovery from heat stress (**Fig. 1, M and N and Movie S1**), demonstrating the ubiquitination is required for stress granule disassembly following heat shock. Similar results were obtained upon siRNA-mediated knockdown of UBA1 (Maxwell et al., co-submitted). Thus, we next sought to investigate the relationship between poly-ubiquitin conjugation of G3BP1 and stress granule dynamics.

### Ubiquitination within G3BP1 NTF2L is required for disassembly of heat shock-induced stress granules

To investigate the effect of G3BP1 ubiquitination on stress granule dynamics, we first sought to identify the heat shock-dependent ubiquitination site(s) within G3BP1. To this end, we generated constructs in which six lysine residues in the NTF2L domain (6KR) or four lysine residues in the RRM domain (4KR) were mutated to arginines. When expressed in *G3BP1/2* dKO cells, G3BP1 6KR showed greatly reduced ubiquitination in response to heat shock, whereas G3BP1 4KR showed no apparent change in poly-ubiquitin conjugation, suggesting that the NTF2L domain is preferentially ubiquitinated upon heat shock (**Fig. 2A**). The NTF2L domain of G3BP1 operates as a dimer, and disruption of dimerization results in stress granule disassembly (*3*). Examination of the crystal structure of the NTF2L dimer (PDB: 5FW5) revealed that all six lysine residues are surface exposed. Four lysine residues (K36, K50, K59, and K64) are clustered to form a positively charged surface on the laterals sides of homodimeric NTF2L domains (**Fig. 2B**), K50 and K76 from each monomer lie adjacent to the dimeric interface, and K123 is located near the hydrophobic grove between alpha helices, where the nsP3 protein of Old World alphaviruses binds to inhibit stress granule assembly (**fig. S2A**). To gain insight into which lysines in G3BP1 are ubiquitinated, we generated a series of K-to-R mutations alone or in combination. Mutation of the four clustered lysine residues (K36/50/59/64R) nearly abolished heat-shock dependent ubiquitination of G3BP1 in *G3BP1/2* dKO cells, whereas other combinations of mutations to NTF2L lysines had no strong effect (**Fig. 2C**). However, ubiquitination levels further decreased in the 6KR mutant, and we therefore used G3BP1 6KR as a ubiquitination-deficient mutant. Notably, ubiquitination of G3BP1 in this region is consistent with results from an independent approach presented in Maxwell et al., co-submitted. Specifically, paired di-Gly and TMT analysis revealed enrichment of ubiquitinated G3BP1 peptide containing K50 (K.NSSYVHGGLDSNGK^50^PADAVYGQK.E) upon heat shock and subsequent decrease in this ubiquitinated species during recovery from stress (**Fig. 2D**). The decrease in ubiquitinated G3BP1 during the recovery phase appears to reflect proteasome-dependent degradation since treatment with the proteasome inhibitor leads to a great accumulation of ubiquitinated G3BP1 (**Fig. 2D**).

**Fig. 2.**
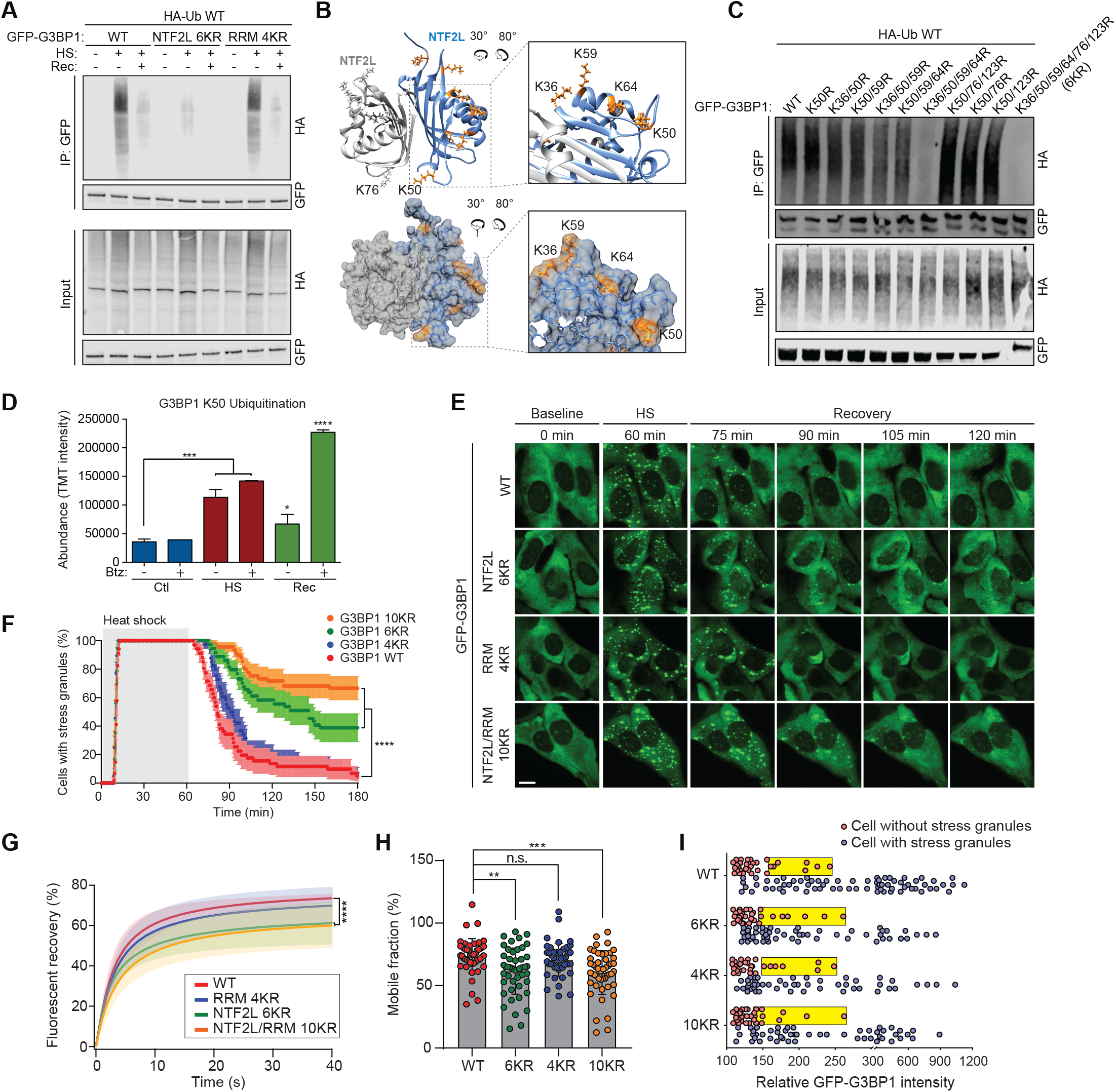
Ubiquitination of the NTF2L domain of G3BP1 is required for the disassembly of stress granules. (**A**) Western blot showing heat shock-induced ubiquitination of G3BP1 NTF2L. HEK293T cells were transfected with HA-Ub and indicated GFP-G3BP1 constructs. Cells were exposed to no stress, heat shock (43°C, 1 h), or heat shock plus recovery (43°C, 1 h; 37°C, 30 min). Cell extracts were captured with magnetic beads conjugated with GFP antibody for IP and resulting beads were analyzed by immunoblot. (**B**) Dimeric structure of G3BP1 NTF2L domain. K36, K50, K59, and K64 are highlighted in enlarged images (orange). (**C**) Western blot showing heat shock-induced ubiquitination of G3BP1 at K36, K50, K59, and K64. HEK293T cells were transfected with HA-Ub and indicated GFP-G3BP1 constructs and exposed to heat shock (43°C, 1 h). Cell extracts were captured with magnetic beads conjugated with GFP antibody for IP and resulting beads were analyzed by immunoblot. (**D**) Peptide abundance (TMT intensity) of ubiquitinated G3BP1 peptides from the ubiquitinome during heat shock (HS) recovery (Rec) in the presence or absence of bortezomib (Btz). Error bars indicate s.d. \**P*= 0.0487, \*\*\**P*= 0.0006 (Ctl vs. HS), \*\*\**P*= 0.0001 (Ctl vs. HS Btz), and \*\*\*\**P*< 0.0001 by one-way ANOVA Dunnett’s multiple comparison test. **(E-F)** Fluorescent imaging of stress granule disassembly in U2OS *G3BP1/2* dKO cells stably expressing G3BP1 WT and 6KR, 4KR, or 10KR mutants. Cells were incubated at 37°C for 2 min, 43°C for 60 min, and 37°C for 118 min, and GFP signals were monitored with 30-sec intervals. Representative still images are shown in (E). The percentage of cells with ≥ 2 stress granules is plotted in (F). Scale bar, 20 μm. Error bars indicate s.e.m. \*\*\*\**P*< 0.0001, Mantel-Cox test. (**G**) FRAP of GFP-G3BP1 within U2OS *G3BP1/2* dKO cells transfected with GFP-G3BP1 WT, 6KR, 4KR, or 10KR, exposed to heat shock (43°C, 1 h) followed by recovery (37°C, 10 min). GFP-positive puncta were analyzed by FRAP beginning after 10 min of recovery. Error bars indicate s.e.m. \**P*< 0.05; \*\*\*\**P*< 0.0001, Tukey’s test. (**H**) Quantification of the percentage of cells with mobile fraction at 40 second recovery. \*\**P*< 0.01; \*\*\**P*< 0.001, by one-way ANOVA Dunnett’s multiple comparison test. **(I)** Intracellular phase diagram of U2OS *G3BP1/2* dKO cells transfected with GFP-G3BP1 WT, 6KR, 4KR, or 10KR and exposed to heat shock (43°C, 1 h). Cells without stress granules (red circles) and with stress granules (blue circles) are plotted according to GFP intensities. Yellow boxes indicate the cells with 25% highest levels of GFP within the stress granule-negative group.

We next sought to determine the role of distinct ubiquitination events in stress granule dynamics, including assembly, disassembly, and dynamic exchange between the stress granule and the surrounding cytoplasm. To this end, we first generated stable cell lines in which G3BP1 6KR, 4KR, or 10KR were re-expressed in *G3BP1/*2 dKO cells at levels comparable to endogenous G3BP1 levels (**fig. S2, B and C**). Upon heat shock, all three mutant forms of G3BP1 assembled stress granules with kinetics identical to WT G3BP1 (**Fig. 2, E and F and Movie S2**), indicating that assembly of stress granules is independent of ubiquitination of G3BP1. This result is consistent with our earlier findings that global inhibition of ubiquitination with TAK-243 did not alter the dynamics of stress granule assembly in response to heat shock (**Fig. 1, M and N**, Maxwell et al., co-submitted). Remarkably, however, 6KR and 10KR mutations resulted in significantly prolonged stress granule disassembly (**Fig. 2, E and F and Movie S2**). G3BP1 10KR (KR mutations in NTF2L and RRM domains) had a more severe impact on stress granule disassembly than G3BP1 6KR (KR mutations in NTF2L only), suggesting that the four lysine residues in the RRM domain may also contribute to stress granule disassembly, although the levels of poly-ubiquitination in the RRM domain were below the detection limit by immunoblotting. FRAP analysis also showed that the G3BP1 6KR and 10KR mutants exhibited a significantly slower recovery rate and a greater immobile fraction in compared with WT and 4KR proteins (**Fig. 2, G and H**), revealing a role for ubiquitination in defining G3BP1 dynamics during recovery from heat shock.

We have previously shown that phase separation of G3BP1 with mRNA triggers stress granule assembly, and, moreover, that the threshold concentration for this phase separation is tuned by intrinsic properties of G3BP1 such as its conformational state and ability to form dimers (*3*). To rule out the possibility that the K-to-R mutations or altered ubiquitination impacts the intrinsic properties of G3BP1 that underlie phase separation, we assessed the impact of KR mutations on dimerization and RNA binding. By a co-immunoprecipitation assay, we confirmed that GFP-G3BP1 WT, 6KR, 4KR, and 10KR mutants all formed dimers with endogenous WT G3BP1 (**fig. S6D**). Moreover, 6KR mutations in the NTF2L domain did not alter G3BP1’s ability to interact with key interacting proteins through this domain (**fig. S2E**), nor did 4KR mutations in the RRM domain alter its ability to bind RNA (**fig. S2F**), suggesting that intrinsic properties of G3BP1 were not compromised by lack of ubiquitination. We also mapped intracellular phase separation thresholds of individual G3BP1 mutants by transiently transfecting G3BP1 into *G3BP1/2* dKO cells and measuring the G3BP1 concentration threshold at which stress granule assembly was initiated (*3*). The concentration threshold for stress granule assembly was similar for G3BP1 WT, 6KR, 4KR, and 10KR mutants (**Fig. 2I**), supporting our interpretation that assembly of stress granules is independent of ubiquitination of G3BP1. Taken together, these results indicate that ubiquitination of G3BP1 is not required for the assembly of stress granules, but selectively impacts their dynamism and is required for their efficient disassembly.

### Heat shock induces interaction of VCP with ubiquitinated G3BP1

A recent study demonstrated that the depletion of cellular ATP by the addition of 2-deoxyglucose (2-DG), an inhibitor of the glycolytic pathway, impairs stress granule assembly and subsequent dynamic behavior, implicating energy-dependent processes in assembly dynamics (*22*). However, the requirement of cellular ATP for the disassembly of stress granules has not been explored. Thus, we sought to examine the contribution of ATP to stress granule disassembly. Under basal conditions, addition of 2-DG led to a ~40% reduction in cellular ATP levels by 30 min as assessed by a luminescent signal-based ATP sensor (**fig. S3A**). In contrast, addition of 2-DG after 60 min heat shock and immediately before recovery led to a ~70% reduction in cellular ATP levels within 10 min (**fig. S3A**). Concurrently, we observed defects in disassembly of stress granules induced by heat shock or arsenite stresses (**fig. S3, B-E, Movie S3**), suggesting that ATP is actively hydrolyzed during recovery processes after heat shock and stress granules are disassembled via energy-dependent processes.

The stress granule proteome contains a large group of proteins with ATPase activities (*22, 23*), and we and others have previously shown that the ATPase valosin-containing protein (VCP) is recruited to stress granules and contributes to their clearance (*12, 15*). VCP is a ubiquitin-dependent protein segregase coupled to the proteasome and autophagy systems (*12, 15, 17*). Since disassembly/clearance of stress granules depends on VCP and the ubiquitination of G3BP1, we hypothesized that VCP acts on ubiquitinated G3BP1 to promote disassembly of stress granules. Thus, we first tested whether recruitment of VCP to stress granules is contingent on ubiquitination of G3BP1. We first confirmed that VCP was recruited to stress granules upon heat shock in WT U2OS cells (**fig. S4A**). Although *G3BP1/2* dKO cells do not form stress granules (*3, 8*), re-expression of GFP-G3BP1 WT or 4KR in these cells restored VCP recruitment to stress granules to levels similar to those observed in WT U2OS cells (**Fig. 3, A and B, fig. S4A**). However, VCP recruitment to stress granules was significantly reduced in *G3BP1/2* dKO cells stably expressing GFP-G3BP1 6KR or 10KR (**Fig. 3, A and B**). This data indicates that ubiquitination of G3BP1 contributes significantly to the recruitment of VCP to stress granules. Whereas G3BP1 is not the only stress granule protein that is ubiquitinated (Maxwell et al., co-submitted), this data is consistent with the fact that G3BP1 is among the most abundant protein constituents in stress granules (*22*)

**Fig. 3.**
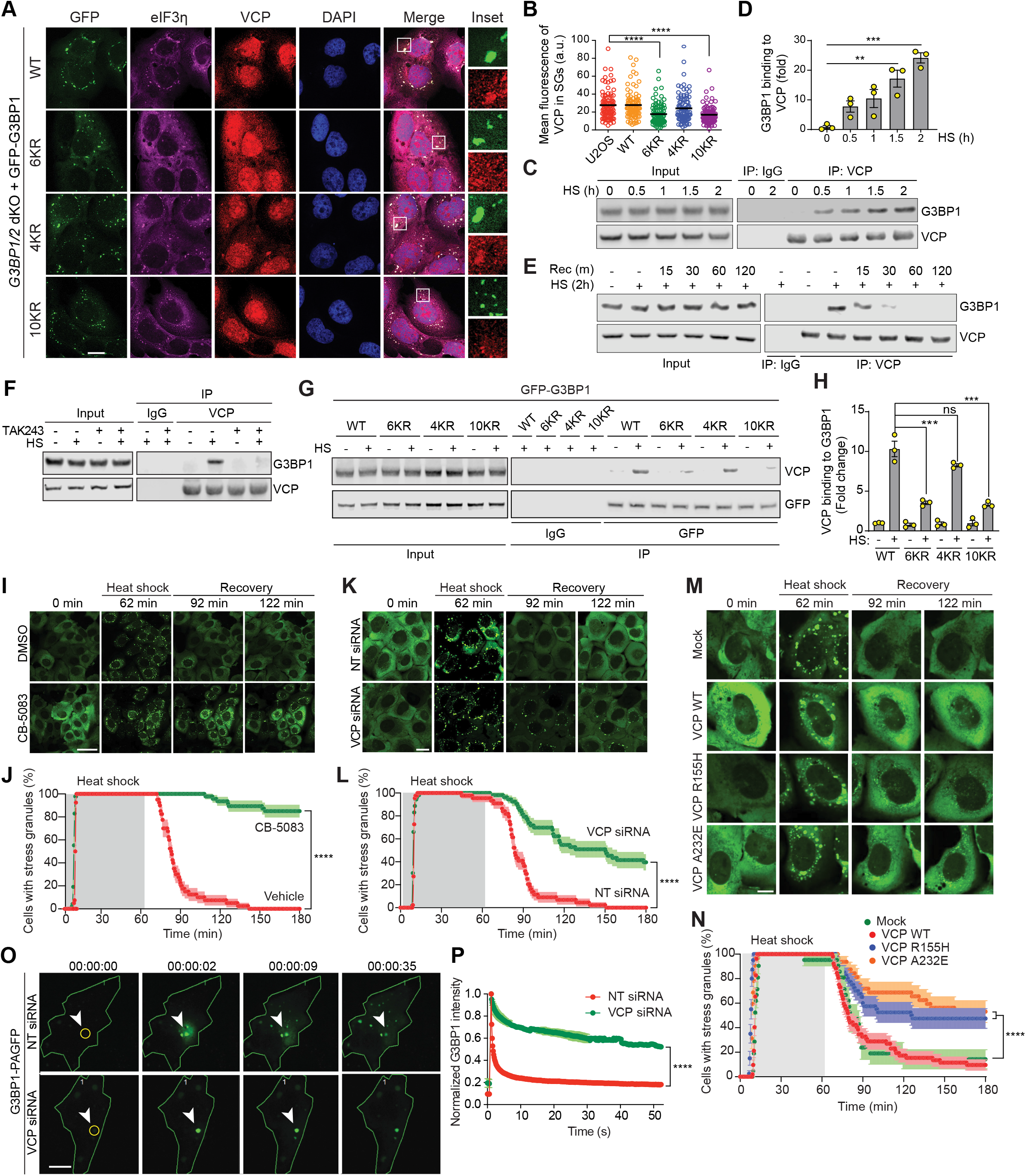
VCP interacts with ubiquitinated G3BP1 in response to heat shock and regulates disassembly of stress granules. **(A-B)** Fluorescent imaging of U2OS *G3BP1/2* dKO cells stably expressing GFP-G3BP WT, 6KR, 4KR or 10KR and exposed to heat shock (43°C, 1 h) (A). Fluorescent intensities of VCP in eIF3η-positive stress granules are plotted in (B). Ubiquitination of G3BP1 is required for recruitment of VCP to stress granules. Scale bar, 20 μm. Error bars indicate s.e.m. \*\*\*\**P*< 0.0001, ANOVA with Tukey’s test. **(C-D)** Western blot of U2OS cell extracts exposed to heat shock (43°C) for indicated times. Cell extracts were captured with magnetic beads conjugated with VCP antibody for IP and resulting beads were analyzed by immunoblot (C). Quantification of immunoblots from 3 replicates are shown in (D). Blots show enhanced interaction between G3BP1 and VCP in response to heat shock. Error bars indicate s.e.m. \*\**P*< 0.01; \*\*\**P*< 0.001, ANOVA with Tukey’s test. **(E)** Western blot showing G3BP1-VCP interaction in U2OS cells based on duration of recovery after heat shock (43°C, 2 h). Cell extracts were captured with magnetic beads conjugated with VCP antibody for IP and resulting beads were analyzed by immunoblot. **(F)** Western blot of U2OS cells treated with DMSO or TAK-243 (60 min) and exposed to heat shock (43°C, 2 h). Cell extracts were captured with magnetic beads conjugated with VCP antibody for IP and resulting beads were analyzed by immunoblot. Blots show an attenuated G3BP1-VCP interaction with inhibition of ubiquitination. **(G-H)** Western blot of U2OS *G3BP1/2* dKO cells stably expressing G3BP1 WT, 6KR, 4KR, or 10KR and exposed to heat shock (43°C, 2 h). Cell extracts were captured with magnetic beads conjugated with GFP antibody for IP and resulting beads were analyzed by immunoblot. Quantification of immunoblots from 3 replicates are shown in (H). Blots show an attenuated interaction between ubiquitination-defective G3BP1 and VCP. Error bars indicate s.e.m. ns, not significant; \*\*\**P*< 0.001, ANOVA with Tukey’s test. **(I-J)** Fluorescent imaging of U2OS cells stably expressing GFP-G3BP1 and treated with DMSO or CB-5083 (1 h) prior to live cell imaging. Cells were incubated at 37°C for 2 min, 43°C for 60 min, and 37°C for 118 min, and GFP signals were monitored with 30-sec intervals. Representative still images are shown in (I). The percentage of cells with ≥ 2 stress granules is plotted in (J). Staining suggests that VCP inhibition disrupts disassembly of heat shock-induced stress granules. Scale bar, 50 μm. Error bars indicate s.e.m. \*\*\*\**P*< 0.0001, Mantel-Cox test. **(K-L)** Fluorescent imaging of U2OS cells stably expressing GFP-G3BP1 and transfected with non-targeting siRNA or VCP siRNA. Cells were incubated at 37°C for 2 min, 43°C for 60 min, and 37°C for 118 min, and GFP signals were monitored with 30-sec intervals. Representative still images are shown in (K). The percentage of cells with ≥ 2 stress granules is plotted in (L). Staining suggests that VCP knockdown delays disassembly of heat shock-induced stress granules. Scale bar, 20 μm. Error bars indicate s.e.m. \*\*\*\**P*< 0.0001, Mantel-Cox test.**(M-N)** Fluorescent imaging of U2OS cells stably expressing GFP-G3BP1 and transfected with empty vector, VCP WT, VCP R155H, or VCP A232E. Cells were incubated at 37°C for 2 min, 43°C for 60 min, and 37°C for 118 min, and GFP signals were monitored with 30-sec intervals. Representative still images are shown in (M). The percentage of cells with ≥ 2 stress granules is plotted in (N). Staining suggests delayed disassembly of stress granules composed of ubiquitination-defective G3BP1. Scale bar, 10 μm (M). Error bars indicate s.e.m. \*\*\*\**P*< 0.0001, Mantel-Cox test. **(O-P)** Imaging of U2Os cells stably expressing photoactivatable G3BP1 (PAGFP-G3BP1) and transfected with either non-targeting siRNA or VCP siRNA. 48 h post transfection, ROIs were activated by 405-nm laser and GFP signals were monitored with 200-ms intervals (O). Graphs show the normalized intensity of GFP signals within ROIs (P). \*\*\*\**P*< 0.0001, two-way ANOVA with Sidak’s multiple comparison test.

Consistent with these findings, endogenous G3BP1 co-immunoprecipitated with VCP upon heat shock (**Fig. 3, C and D**). This interaction gradually increased during a 2-hour heat shock period and decreased to baseline levels within one hour of recovery, consistent with the progressive, proteasome-dependent drop in levels of ubiquitinated G3BP1 (**Fig. 3C-E**). Interestingly, TAK-243 treatment abolished the VCP-G3BP1 interaction upon heat shock (**Fig. 3F**), suggesting that heat shock induces ubiquitination-dependent interaction of VCP and G3BP1. To further assess the role of G3BP1 ubiquitination in this interaction, we expressed WT, 6KR, 4KR, and 10KR mutants in *G3BP1/2* dKO cells and assessed their binding to VCP. Of note, 6KR and 10KR mutants showed significantly reduced binding to VCP compared to WT G3BP1 upon heat shock, whereas VCP interaction was not significantly impacted in the 4KR mutant (**Fig. 3, G and H**). This observation is consistent with a role for ubiquitination of the NTF2L domain, but not the RRM domain, G3BP1 as primarily responsible for interaction of G3BP1 with VCP. Both 6KR and 10KR mutants still showed modest interaction with VCP upon heat shock, although the levels were substantially lower than WT. Since we did not observe VCP-G3BP1 interaction in cells treated with TAK-243, this finding may be attributable to indirect interaction of VCP with other ubiquitinated proteins bound to G3BP1. However, we cannot eliminate the possibility that either G3BP1 ubiquitination occurs beyond these 10 lysine residues or that the VCP-G3BP1 interaction may occur at low levels independently of the ubiquitination status of G3BP1. Finally, we observed that the levels of ubiquitinated G3BP1 were increased by the addition of a VCP inhibitor (CB-5083) and stabilized by the addition of a proteasome inhibitor (bortezomib), suggesting that ubiquitinated G3BP1 may be targeted by VCP for proteasomal degradation upon recovery (**fig. S4, B and C**).

### VCP is required for disassembly of stress granules

VCP has been implicated in multiple routes of VCP elimination, including clearance of stress granules by autophagy (*6, 11, 12*) and disassembly of stress granules to enable recycling of constituents (*17*). Heat shock-dependent ubiquitination of G3BP1 and subsequent interaction with VCP prompted us to determine whether VCP activity is required for disassembly of heat shock-induced stress granules. We observed that either chemical inhibition of VCP using two different inhibitors (**Fig. 3, I and J, fig. S4, D and E, and Movie S4**) or siRNA knockdown of VCP (**Fig. 3, K and L and Movie S5**) led to significantly delay in the disassembly of heat shock-induced stress granules. We have previously observed that expression of disease-causing mutant forms of VCP (A232E and R155H) causes the accumulation of spontaneously arising, poorly dynamic stress granules that recruit TDP-43 (*12*). Similar TDP-43-laden, spontaneously arising, poorly dynamic stress granules are found in cells expressing disease mutant forms of RNA-binding proteins, such as FUS, hnRNPA1, hnRNPA2 and TIA-1 (*11*). These prior observations suggested the possibility that mutations in VCP impair its ability to disassemble stress granules, representing a mechanistic intersection for multiple, distinct disease-causing mutations. Thus, we tested the impact of mutant VCP on stress granule dynamics. Whereas expression of WT VCP did not alter the dynamics of stress granules, expression of VCP A232E or R155H significantly delayed disassembly of stress granules upon removal of heat stress (**Fig. 3, M and N and Movie S6**). This result suggests that mutant VCP proteins have a dominant negative effect on stress granule disassembly similar to that observed in the presence of VCP inhibitors or VCP knockdown.

To further assess the impact of VCP on G3BP1 dynamics in stress granules, we generated a stable U2OS cell line expressing G3BP1 conjugated to photoactivable GFP (G3BP1-PAGFP) (*24*) that allowed us to explore intracellular G3BP1 dynamics by tracking photoactivated molecules. Regions of interest (ROIs) within cells expressing G3BP1-PAGFP were excited briefly with a ~400-nm laser to activate selected pools of G3BP1, followed by time-lapse 488-nm imaging. Stress granule-localized G3BP1 proteins were successfully visualized after PAGFP photoconversion (**Fig. 3O**). In cells transfected with non-targeting siRNA, the fluorescence intensity of stress granules diminished over time within ROIs and redistributed to nearby stress granules (**Fig. 3, O and P**). Remarkably, photoactivated G3BP1-PAGFP in cells depleted of VCP displayed significantly longer residence time in stress granules and limited redistribution of signal to neighboring stress granules, suggesting that VCP is required for unloading G3BP1 from stress granules (**Fig. 3, O and P, fig. S4F, and Movies S7 and S8**).

### FAF2, an ER-resident VCP adaptor, links ER to G3BP1 for disassembly of stress granules

VCP associates with a large number of interacting partners and cofactors that regulate its activities in a variety of cellular pathways (*25–28*). Although VCP is known to have some affinity for ubiquitin, it binds to ubiquitinated substrates largely through cofactor proteins with ubiquitin-binding domains that function as ubiquitin adaptors for VCP (*29*). Approximately 35 VCP cofactor proteins have been identified in mammalian cells (*15, 28, 30, 31*), although in most cases the pathways in which these cofactors function, and the substrates that they recognize, remain poorly defined. Thus, we sought to identify the cofactor(s) that link VCP to stress granules by binding ubiquitinated G3BP1 and thereby influencing disassembly of stress granules.

To establish a candidate list of relevant VCP interactors, we integrated a list of 35 VCP cofactors with 1,552 proteins previously identified in stress granules initiated by multiple stressors (**Table 1**) (*3, 22, 32, 33*). The stress granule proteome and VCP cofactors had 7 proteins in common (**Fig. 4A**), including ZFAND1, a protein that has been described to promote the clearance of arsenite-induced stress granules by recruiting VCP and the 26S proteasome (*15*). We confirmed interaction of 5 of these adaptors with endogenous VCP (**fig. S5A**). Interestingly, of the 7 proteins in common between the stress granule proteome and known VCP cofactors, only FAS-associated factor 2 (FAF2) demonstrated interaction with G3BP1 upon heat shock, suggesting the possibility that FAF2 binding to VCP might mediate the interaction between VCP and G3BP1 (**Fig. 4B**). To test whether FAF2 is an adaptor protein that links VCP to G3BP1, we assessed VCP interaction with G3BP1 in the presence or absence of FAF2 and found that the VCP-G3BP1 interaction was strictly dependent on FAF2 (**Fig. 4C and fig. S5B**). The FAF2-G3BP1 interaction was G3BP1 ubiquitination-dependent, as demonstrated by a loss of interaction of FAF2 with G3BP1 6KR and 10KR upon heat shock (**Fig. 4D**). Furthermore, depletion of FAF2 by siRNA delayed disassembly of heat shock-induced stress granules upon removal of heat stress, phenocopying our earlier observations using chemical inhibitors of VCP (**Fig. 4, E and F, and Movie S9**). In contrast, depletion of 6 other adaptors did not significantly alter the disassembly of heat shock-induced stress granules upon removal of heat stress (**fig. S5C**), suggesting that VCP specifically engages FAF2 to regulate dynamics of stress granules. Of note, depletion of ZFAND1 delayed disassembly of arsenite-induced stress granules upon removal of arsenite as previously demonstrated (*15*), whereas depletion of FAF2 fail to delay disassembly of arsenite-induce stress granules, demonstrating a context-dependent control of stress granule dynamics by ZFAND1 and FAF2 (**fig. S5, D-G**).

**Fig. 4.**
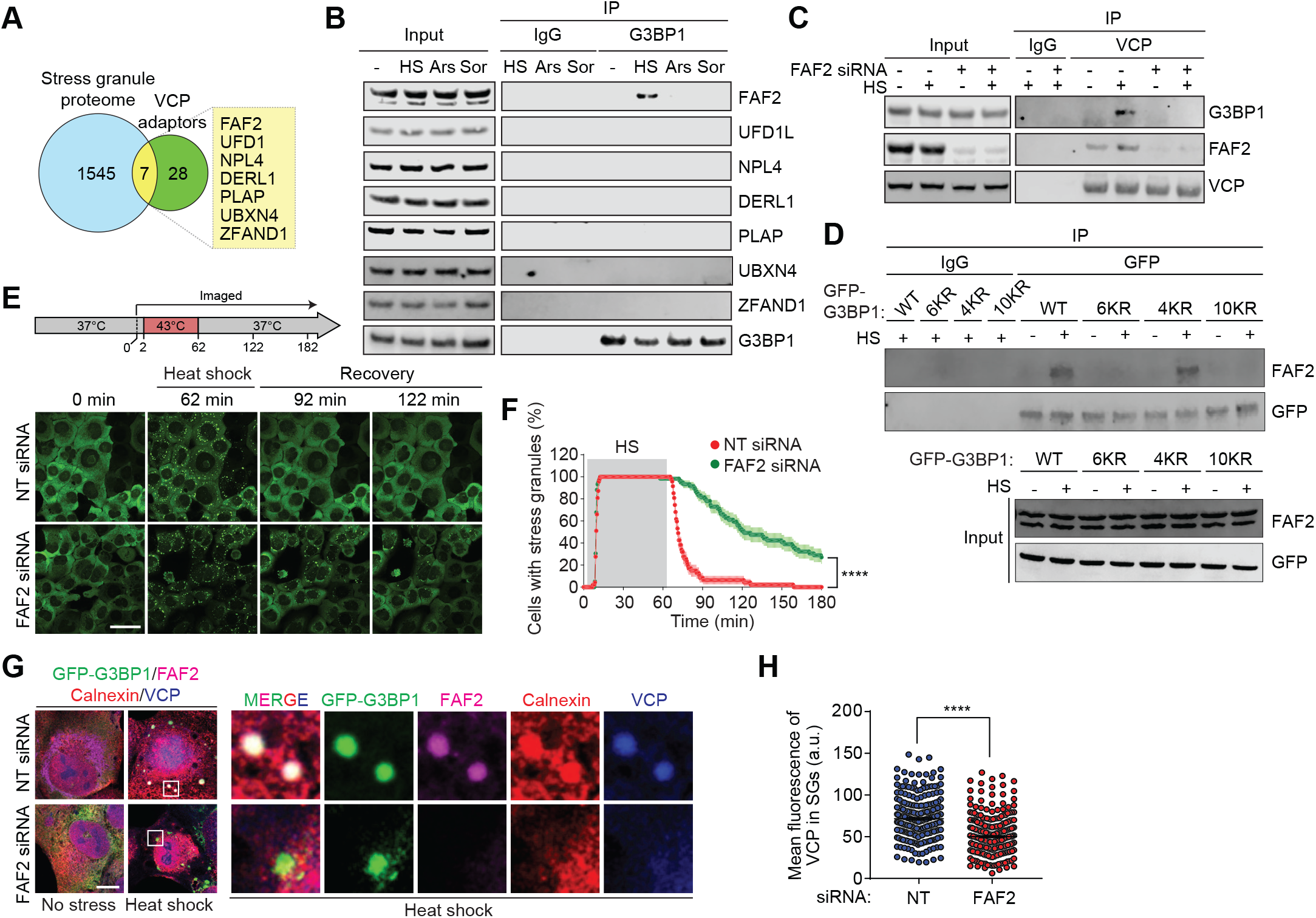
FAF2 links ubiquitinated G3BP1 to VCP. **(A)** Venn diagram showing overlapping proteins among the stress granule proteome and known VCP adaptors. **(B)** Western blot of U2OS cells exposed to no stress, heat shock (43°C, 1.5 h), oxidative stress (0.5 mM sodium arsenite, 1.5 h), or osmotic stress (0.4 M sorbitol, 1.5 h). Cell extracts were captured with magnetic beads conjugated with G3BP1 antibody for IP and resulting beads were analyzed by immunoblot. Blots show interaction between G3BP1 and FAF2 upon heat shock. **(C)** Western blot of U2OS cells transfected with non-targeting siRNA or FAF2 siRNA and exposed to heat shock (43°C, 1.5 h). Cell extracts were captured with magnetic beads conjugated with VCP antibody for IP and resulting beads were analyzed by immunoblot. Blots suggest a requirement of FAF2 for interaction between G3BP1 and VCP. (D) Western blots of U2OS *G3BP1/2* dKO cells stably expressing G3BP1 WT, 6KR, 4KR or 10KR mutants and exposed to heat shock (43°C, 2 h). Cell extracts were captured with magnetic beads conjugated with GFP antibody for IP and resulting beads were analyzed by immunoblot. Blots suggest impaired interaction between ubiquitination-defective G3BP1 and FAF2. **(E-F)** Fluorescent imaging of U2OS cells stably expressing GFP-G3BP1 and transfected with non-targeting siRNA or FAF2 siRNA. Cells were incubated at 37°C for 2 min, 43°C for 60 min, and 37°C for 118 min, and GFP signals were monitored with 30-sec intervals. Representative still images are shown in (E). The percentage of cells with ≥ 2 stress granules is plotted in (F). Imaging shows delayed disassembly of stress granules by FAF2 knockdown. Scale bar, 50 μm. Error bars indicate s.e.m. \*\*\*\**P*< 0.0001, Mantel-Cox test. **(G-H)** Fluorescent imaging of U2OS cells stably expressing GFP-G3BP1 and exposed heat shock (43°C, 60 min). Fluorescent intensities of VCP in stress granules are plotted in (H). Imaging shows recruitment of VCP- and FAF2-resident stress granules to ER (calnexin) upon heat shock. Scale bar, 10 μm. Error bars indicate s.e.m. \*\*\*\**P*< 0.0001, ANOVA with Tukey’s test.

FAF2, also known as UBXD8, is an ER-resident protein that is best known for mediating ubiquitin-dependent degradation of misfolded ER proteins (*34, 35*). Intriguingly, the VCP-FAF2 complex has been shown to disassemble mRNPs by promoting the release of ubiquitinated HuR, an RNA-binding protein, from mRNA (*36*). Furthermore, a recent study described ER tubules as platforms for fission of membraneless organelles, including processing bodies and stress granules (*18*), providing an unanticipated role for the ER in regulating the assembly and disassembly of membraneless organelles. Thus, we asked whether FAF2 connects the ER to stress granules by interacting with ubiquitinated G3BP1.

We began by examining the spatiotemporal relationship between the ER, FAF2, G3BP1, and VCP upon heat shock. At baseline, FAF2 was found diffusely distributed with the ER, as indicated by colocalization with the ER marker calnexin (**fig. S6A**). Upon heat shock, calnexin and FAF2 signals was found in protrusions from ER that colocalized with stress granules marked by GFP-G3BP1 (**Fig. 4G**). VCP showed a similar colocalization with G3BP1, FAF2 and ER marker upon heat shock (**Fig. 4G**). In contrast, arsenite stress did not induce FAF2 localization with stress granules (**fig. S6B**). Notably, VCP accumulation in heat stress-induced stress granules was significantly decreased in cells depleted of FAF2, whereas the more modest VCP accumulation in arsenite-induced stress granules was not influenced by depletion of FAF2 (**Fig. 4H and fig. S6, C and D**), supporting our hypothesis that FAF2 functions as a VCP adaptor that connects the ER to heat shock-induced stress granules by interacting with ubiquitinated G3BP1.

## Final Comments

This study presents a striking example of customize, context-dependent cellular response to stress. Whereas stress granule assembly is evidently a universal process across cell types and in response to a wide variety of stresses, it has recently become apparent that the composition of stress granules is context-dependent with respect to cell type and the initiating stress (*32*). Our study builds on this perspective by demonstrating context-dependent differences in stress granule disassembly. This context is particularly important when considering how we investigate the relationship of stress granules to disease: we must be attentive to disease-relevant contexts, including the use of specific cell types or the administration of different types of stress.

Much remains to be learned about the distinct mechanisms of stress granule disassembly, and in particular how these mechanisms may be impacted by disease-causing mutations. Indeed, although we previously showed that disease mutations in VCP impair autophagy-dependent stress granule clearance (*12*), here we found that these mutations also impair autophagy-independent disassembly. Furthermore, our results demonstrate that disassembly is mediated by extraction of ubiquitinated G3BP, a finding that underscores the central role for G3BP in maintaining the stress granule interaction network. Moreover, informed by the observation by Maxwell et al. that cells produce distinct patterns of ubiquitination in response to different stressors, our study illustrates how even a single protein within a stress-specific ubiquitinome can be pursued experimentally to demonstrate meaningful consequences for cellular function.

## Acknowledgements

We thank Natalia Nedelsky for editorial assistance. We thank the Center for Advanced Genome Engineering at St. Jude Children’s Research Hospital for assistance with CRISPR-Cas9 modified cell lines, as well as J. Messing (St. Jude) and J. Temirov (St. Jude) for assistance with microscopy.

## Author contributions

Y.G., H.J.K., and J.P.T. conceived the project and J.P.T. supervised the project. Y.G. performed all experiments except G3BP1 di-GLY and TMT analysis performed by B.A.M., G3BP1-PAGFP photoactivation, G3BP1-GFP live-cell imaging upon Eerl treatment performed by R.M.K, and stress granule core proteome analysis by P.Z. Data was analyzed by Y.G., B.A.M., R.M.K., H.J.K., and J.P.T. H.J.K., and J.P.T. wrote the primary draft of manuscript and all authors contributed to the final version.

## Data and materials availability

All data is available in the main text or the supplementary materials.

## Supplementary Materials

Materials and Methods

Figures S1-S6

Movies S1-S9

Table 1

References

## References and Notes

1. J. R. Buchan, mRNP granules. Assembly, function, and connections with disease. RNA Biol 11, 1019–1030 (2014).

2. E. Gomes, J. Shorter, The molecular language of membraneless organelles. J Biol Chem294, 7115–7127 (2019).

3. P. Yang et al., G3BP1 Is a Tunable Switch that Triggers Phase Separation to Assemble Stress Granules. Cell 181, 325–345 e328 (2020).

4. D. W. Sanders, et al., Competing Protein-RNA Interaction Networks Control Multiphase Intracellular Organization. Cell 181, 306–324 e328 (2020).

5. J. Guillen-Boixet et al., RNA-Induced Conformational Switching and Clustering of G3BP Drive Stress Granule Assembly by Condensation. Cell 181, 346–361 e317 (2020).

6. C. Mathieu, R. V. Pappu, J. P. Taylor, Beyond aggregation: Pathological phase transitions in neurodegenerative disease. Science 370, 56–60 (2020).

7. S. F. Banani, H. O. Lee, A. A. Hyman, M. K. Rosen, Biomolecular condensates: organizers of cellular biochemistry. Nat Rev Mol Cell Biol 18, 285–298 (2017).

8. N. Kedersha et al., G3BP-Caprin1-USP10 complexes mediate stress granule condensation and associate with 40S subunits. J Cell Biol 212, 845–860 (2016).

9. P. Zhang et al., OptoGranules reveal the evolution of stress granules to ALS-FTD pathology. bioRxiv (2018).

10. D. Tauber, G. Tauber, R. Parker, Mechanisms and Regulation of RNA Condensation in RNP Granule Formation. Trends in biochemical sciences 45, 764–778 (2020).

11. N. B. Nedelsky, J. P. Taylor, Bridging biophysics and neurology: aberrant phase transitions in neurodegenerative disease. Nat Rev Neurol 15, 272–286 (2019).

12. J. R. Buchan, R. M. Kolaitis, J. P. Taylor, R. Parker, Eukaryotic stress granules are cleared by autophagy and Cdc48/VCP function. Cell 153, 1461–1474 (2013).

13. P. Anderson, N. Kedersha, RNA granules: post-transcriptional and epigenetic modulators of gene expression. Nat Rev Mol Cell Biol 10, 430–436 (2009).

14. S. Markmiller et al., Active Protein Neddylation or Ubiquitylation Is Dispensable for Stress Granule Dynamics. Cell Rep 27, 1356–1363 e1353 (2019).

15. A. Turakhiya et al., ZFAND1 Recruits p97 and the 26S Proteasome to Promote the Clearance of Arsenite-Induced Stress Granules. Mol Cell 70, 906–919 e907 (2018).

16. J. van den Boom, H. Meyer, VCP/p97-Mediated Unfolding as a Principle in Protein Homeostasis and Signaling. Mol Cell 69, 182–194 (2018).

17. B. Wang et al., ULK1 and ULK2 Regulate Stress Granule Disassembly Through Phosphorylation and Activation of VCP/p97. Mol Cell 74, 742–757 e748 (2019).

18. J. E. Lee, P. I. Cathey, H. Wu, R. Parker, G. K. Voeltz, Endoplasmic reticulum contact sites regulate the dynamics of membraneless organelles. Science 367, (2020).

19. K. N. Swatek, D. Komander, Ubiquitin modifications. Cell Res 26, 399–422 (2016).

20. F. Ohtake, Y. Saeki, S. Ishido, J. Kanno, K. Tanaka, The K48-K63 Branched Ubiquitin Chain Regulates NF-kappaB Signaling. Mol Cell 64, 251–266 (2016).

21. P. V. Hornbeck, et al., PhosphoSitePlus, 2014: mutations, PTMs and recalibrations. Nucleic Acids Res 43, D512–520 (2015).

22. S. Jain et al., ATPase-Modulated Stress Granules Contain a Diverse Proteome and Substructure. Cell 164, 487–498 (2016).

23. B. D. Freibaum, J. Messing, P. Yang, H. J. Kim, J. P. Taylor, High fidelity reconstitution of stress granules and nucleoli in mammalian cellular lysate. bioRxiv, (2020).

24. G. H. Patterson, J. Lippincott-Schwartz, A photoactivatable GFP for selective photolabeling of proteins and cells. Science 297, 1873–1877 (2002).

25. H. Meyer, M. Bug, S. Bremer, Emerging functions of the VCP/p97 AAA-ATPase in the ubiquitin system. Nat Cell Biol 14, 117–123 (2012).

26. A. Stolz, W. Hilt, A. Buchberger, D. H. Wolf, Cdc48: a power machine in protein degradation. Trends in biochemical sciences 36, 515–523 (2011).

27. H. O. Yeung, et al., Insights into adaptor binding to the AAA protein p97. Biochem Soc Trans 36, 62–67 (2008).

28. Y. Ye, W. K. Tang, T. Zhang, D. Xia, A Mighty “Protein Extractor” of the Cell: Structure and Function of the p97/CDC48 ATPase. Front Mol Biosci 4, 39 (2017).

29. Y. Ye, Diverse functions with a common regulator: ubiquitin takes command of an AAA ATPase. J Struct Biol 156, 29–40 (2006).

30. H. Meyer, C. C. Weihl, The VCP/p97 system at a glance: connecting cellular function to disease pathogenesis. J Cell Sci 127, 3877–3883 (2014).

31. D. Burana et al., The Ankrd13 Family of Ubiquitin-interacting Motif-bearing Proteins Regulates Valosin-containing Protein/p97 Protein-mediated Lysosomal Trafficking of Caveolin 1. J Biol Chem 291, 6218–6231 (2016).

32. S. Markmiller et al., Context-Dependent and Disease-Specific Diversity in Protein Interactions within Stress Granules. Cell 172, 590–604 e513 (2018).

33. J. Y. Youn, et al., High-Density Proximity Mapping Reveals the Subcellular Organization of mRNA-Associated Granules and Bodies. Mol Cell 69, 517–532 e511 (2018).

34. B. Mueller, E. J. Klemm, E. Spooner, J. H. Claessen, H. L. Ploegh, SEL1L nucleates a protein complex required for dislocation of misfolded glycoproteins. Proc Natl Acad Sci U S A 105, 12325–12330 (2008).

35. Y. Xia et al., Pathogenic mutation of UBQLN2 impairs its interaction with UBXD8 and disrupts endoplasmic reticulum-associated protein degradation. J Neurochem 129, 99–106 (2014).

36. H. L. Zhou, C. Geng, G. Luo, H. Lou, The p97-UBXD8 complex destabilizes mRNA by promoting release of ubiquitinated HuR from mRNP. Genes Dev 27, 1046–1058 (2013).

