## Supplemental text and figures for "Ubiquitination of G3BP1 mediates stress granule disassembly in a stress-specific manner"

**This PDF file includes:**

Materials and Methods
Figs. S1 to S6

**Other Supplementary Materials for this manuscript include the following:**

Table S1
Movies S1 to S9

#### Materials and Methods

Plasmid, siRNA and transfection

Synthetic G3BP1 fragments with K36/50/59/64/76/123R (NTF2L 6KR), K353/357/376/393R (RRM 4KR), and K36/50/59/64/76/123/353/357/376/393R (NTF2L/RRM 10KR) mutations were inserted into *HindIII* and *BamHI* sites of pEGFP-C3 (Clontech) by Bio Basic Inc. pEGFP-C3 G3BP1 K50R, K36/50R, K50/59R, K36/50/59R, K50/59/64R, K36/50/59/64R, K50/76R, K50/123R, and K50/76/123R mutants were generated by site-directed mutagenesis using a Q5 Site-Directed Mutagenesis kit (New England Biolabs; E0054S). pEGFP-C3 G3BP1 ΔIDR1/2 constructs have been previously described (*1*). pRK5-HA-ubiquitin-WT, pRK5-HA-ubiquitin-K48R, pRK5-HA-ubiquitin-K63R, pRK5-HA-ubiquitin-K48 and pRK5-HA-ubiquitin-K63 (Addgene; 17608, 17604, 17606, 17605, and 17606, respectively) were kindly provided by Dr. Ted Dawson. pcDNA3.1-FLAG-VCP WT, VCP R155H, and VCP A232E have been previously described (*2*). pCLi40w-MND-G3BP1-GFP and pCLiw40-MND-G3BP1-PAGFP plasmids, used to generate G3BP-GFP stable U2Os cells via lentiviral transduction, were constructed by releasing dsRedEX2-EIF1α-GFP from pCLEG-MND-dsRedEX2 (Vector Development and Production, SJCRH) and inserting G3BP1-GFP or G3BP1-PAGFP into the *EcoRI* and *BsrGI* positions. G3BP1-GFP and G3BP1-PAGFP inserts were released from peGFP-N1-G3BP and pePAGFP-N1-G3BP by *EcoRI* and *dBsrGI* digestion. G3BP1 was inserted in peGFP-N1 or pePAGFP-N1 (Clontech) with primers containing *EcoRI* and *BamHI* sites in the 5′ and 3′ sites of G3BP. Similarly, the pCLi40w-MND-TIAL1-PAGFP construct was generated using *EcoRI* and *KpnI*. For knockdown experiments, the following siRNA constructs were purchased from Horizon Discovery: pooled nontargeting siRNA (D-001810-10), VCP (L-008728-00), FAF2 (L-010649-02), UFD1L (L-017918-00), NPL4 (L-020796-01), DERL1 (L-010733-02), PLAP (L-016215-00), UBXN4 (L-014184-01) and ZFAND1 (L-009638-02). Transporter 5 (Polysciences; 26008) and FuGENE 6 (Promega; E2691) were used for transient transfections of cDNA into HEK293T and U2OS cells, respectively, according to the manufacturer's instructions. Lipofectamine RNAiMax (Thermo Fisher Scientific; 13778150) was used for transfection of siRNA according to the manufacturer's instructions.

Cell culture

HEK293T (CRL-3216) and U2OS (HTB-96) cells were purchased from ATCC, cultured in Dulbecco's modified Eagle's medium (HyClone) supplemented with 10% fetal bovine serum (HyClone; SH30396.03), 1X GlutaMAX (Thermo Fisher Scientific; 35050061), 50 U/ml penicillin-50 μg/ml streptomycin (Gibco; 15140–122) and maintained at 37°C in a humidified incubator with 5% $CO_2$. U2OS *G3BP1/2* KO cells have been previously described (*3*). U2OS cells stably expressing GFP-G3BP1 have been previously described (*4*).

Generation of G3BP1 add-back cell lines

U2OS *G3BP1/2* dKO cells were transfected with pEGFP-C3 G3BP1 WT, NTF2L 6KR, RRM 4KR, or NTF2L/RRM 10KR constructs using FUGENE 6 (Promega). 48 h after transfection, 500 μg/ml G418 sulfate (Thermo Fisher Scientific; 10131035) was added to culture media without penicillin and streptomycin for selection. After pharmacological selection, GFP-positive cells were purified using cell sorting to produce stable cell lines.

Heat shock and drug treatments

For heat shock, cells were transferred to a 43°C humidified incubator with 5% $CO_2$. Chemicals dissolved in DMSO were prepared and added to cells at the following concentrations: sodium arsenite (0.5 mM, Millipore Sigma; 1062771000), TAK-243 (1 μM, ChemieTek; CT-M7243), CB-5083 (1 μM, Cayman Chemical; 19311) and bortezomib (1 μM, Millipore Sigma; 5043140001). D-Sorbitol (0.4 M, Millipore Sigma; S1876) was dissolved in culture media and pre-warmed before treatment. Eeyarestatin I was used at a final concentration of 56 μM (30 min pretreatment).

Western blotting

Cells were washed twice with PBS and lysed with RIPA buffer (25 mM Tris-HCl, pH 7.6, 150 mM NaCl, 1% NP-40, 1% sodium deoxycholate, 0.1% SDS; Thermo Scientific; 89901) supplemented with 1 mM EDTA (Invitrogen; 15575020) and proteinase inhibitor cocktail (Roche; 1183617001). Lysates were centrifuged for 15 min at 4°C at 20,000 x g. 4X NuPAGE LDS sample buffer (Thermo Fisher Scientific; NP0007) was added to the supernatant and samples were boiled at 90°C for 5 min. Samples were separated in 4%–12% NuPAGE Bis-Tris gels (Thermo Fisher Scientific; NP0336BOX or NP0321BOX) and transferred to PVDF membranes (Thermo Fisher Scientific; IB24001) using an iBlot 2 transfer device (Thermo Fisher Scientific). Membranes were blocked with Odyssey blocking buffer (LI-COR Biosciences; 927-50000) and then incubated with primary antibodies at 4°C overnight: G3BP1 (Proteintech; 13057-2-AP), ubiquitin (Santa Cruz Biotechnology; sc-8017), HA tag (Invitrogen; 715500), GFP (Invitrogen; A11122), VCP (Santa Cruz Biotechnology; sc-20799 or Thermoscientific; MA3-004), FAF2 (Proteintech; 16251-1-AP), UFD1L (Proteintech; 10615-1-AP), NPL4 (Proteintech; 11638-1-AP), DERL1 (Thermo Scientific; PA553444), PLAP (Santa Cruz Biotechnology; sc-390454), UBXN4 (Thermo Fisher Scientific; PA5577611), ZFAND1 (Sigma-Aldrich; HPA023383), β-actin (Santa Cruz Biotechnology; sc-47778 or Sigma-Aldrich; A5316), and GAPDH (Santa Cruz Biotechnology; sc-32233 or Sigma-Aldrich; G9545). Membranes were washed 3 times with TBS-T (0.05% Tween) and further incubated with IRDye 680RD/800CW-labeled secondary antibodies (LI-COR Biosciences; 926-68073 or 926-32212) at a dilution of 1:10,000. Membranes were visualized with an Odyssey Fc imaging system (LI-COR Biosciences) and quantified using ImageJ software (NIH).

Immunoprecipitation

Cells were washed twice with PBS and lysed with IP lysis buffer (25 mM Tris-HCl, pH 7.4, 150 mM NaCl, 1% NP-40, 1 mM EDTA, 5% glycerol; Thermo Scientific; 87787) supplemented with 20 mM N-ethylmaleimide (NEM) (Sigma-Aldrich; E3876), 50 μM PR-619 (Sigma-Aldrich, 662141) and proteinase inhibitor cocktail (Roche; 1183617001). Lysates were centrifuged at 4°C for 15 min at 20,000 x g. The supernatants were incubated with normal mouse IgG (Santa Cruz Biotechnology; sc-2025), GFP antibody (Santa Cruz Biotechnology; sc-9996), VCP antibody (Santa Cruz Biotechnology; sc-57492), or G3BP1 antibody (BD; 611127) conjugated with protein A/G magnetic beads (Thermo Fisher Scientific; 88803) at 4°C overnight. Beads were washed three times with wash buffer (50 mM Tris-HCl, pH 7.5, 500 mM NaCl, 0.05% Tween-20, 20 mM NEM and 50 μM PR-619) and treated with 0.1 M glycine pH 2.7 (Teknova; G4527) at room temperature for 10 min to elute proteins. The resulting samples were analyzed by Western blotting.

TUBE pulldown

Lysates were prepared using the same procedures as when performing immunoprecipitation. Lysates were incubated with Halo-beads or Halo-4xUBA$^{UBQLN1}$-beads at 4°C overnight. Beads were washed three times with wash buffer, mixed with 4X NuPAGE LDS sample buffer, and boiled at 90°C for 5 min. The resulting samples were analyzed by Western blotting.

Immunofluorescence

Cells were grown in 8-well chamber slides (Millipore; PEZGS0816). Cells were fixed with 4% paraformaldehyde (Alfa Aesar; J61899) in PBS for 10 min, permeabilized with 0.2% Triton-X100 in PBS for 5 min, and blocked with 3% BSA for 1 h. Samples were further incubated with primary antibodies as the following targets in blocking buffer at 4°C overnight: Lys48-linkage specific ubiquitin (Millipore Sigma; 05-1307), Lys63-linkage specific ubiquitin (Millipore Sigma; 05-1308), G3BP1 (BD; 611127), G3BP1 (Proteintech; 13057-2-AP), eIF3η (Santa Cruz Biotechnology; sc-16377), VCP (BD Biosciences 612183), FAF2 (Proteintech; 16251-1-AP), and calnexin (Thermo Fisher Scientific; PA5-19169). Samples were washed 3 times with PBS and incubated with host-specific Alexa Fluor 488/555/647 secondary antibodies (Thermo Fisher Scientific) for 1 h at room temperature. For microscopic imaging, slides were mounted with ProLong Gold Antifade reagent with DAPI (Thermo Fisher Scientific; P36931). Images were captured using a Leica TCS SP8 STED 3X confocal microscope with a 63x oil objective. To stain ER-resident calnexin, cells were grown in fibronectin-coated coverslips (neuVitro; GG18FIBRONECTIN) and permeabilized with 0.2% Triton-X100 in PBS for 1 min. Slides were mounted with ProLong Glass Antifade reagent (Thermo Fisher Scientific; P36984).

Disassembly of stress granules, intracellular phase diagram, and FRAP using time-lapse live-cell microscopy

A Yokogawa CSU W1 spinning disk attached to a Nikon Ti2 eclipse with a Photometrics Prime 95B camera using Nikon Elements software was used in time-lapse live-cell imaging and FRAP. The light path was split between the port for the spinning disk/acquisition laser and the FRAP lasers, enabling FRAP to occur simultaneously while imaging. Imaging was taken using a 60x Plan Apo 1.4NA oil objective and Perfect Focus 2.0 (Nikon) engaged the duration of the capture. During imaging, cells were maintained at 37°C and supplied with 5% $CO_2$ using a Bold Line Cage Incubator (Okolabs) and an objective heater (Biotechs).

To monitor the disassembly of heat shock-induced stress granules, multipoint images over 5 xy fields for each condition per one replicate were taken with the 555-nm laser. At 2 min into imaging, the objective temperatures were raised to 43°C for 60 min. After heat shock, the temperature was lowered back to 37°C to alleviate the stress, and cells were imaged after 2 h had passed. Images were taken at each xy position every 30 s.

For intracellular phase diagrams, multipoint images over 25 xy fields for each condition per one replicate were taken with the 555-nm laser. At 2 min into imaging, the objective temperatures were raised to 43°C for 60 min. Images were taken at each xy position every 1 min. Phase diagrams were constructed by measuring GFP fluorescence intensity in each cell and assessing the presence of SGs using Fiji software.

For fluorescence recovery after photobleaching, time lapses were acquired every 100 ms over the course of 45 sec for stress granules with photobleaching with the 488-nm FRAP laser occurring 2 sec into capture. Data were taken from at least n =10 different cells or lysate granules for each condition. In Nikon Elements, ROIs were generated in the photobleached region, a non-photobleached cell, and the background for each time lapse, and the mean intensity of each was extracted. For photobleached regions, a 2.5 μm-diameter circle was used. Data was repeated in triplicate for each condition, with each replicate having at least n = 10 cells. The values retrieved from the ROIs were exported into Igor Pro 7.0 (WaveMetrics) and fit curves were generated after photobleach and background values were corrected.

Photoactivation of G3BP1

For live imaging of photoactivatable GFP, U2OS cells expressing G3BP-GFP or G3BP1-PAGFP were plated on 40-mm #1.5 thick coverslips (Bioptechs) or chambered coverglass (Millipore). Cells were observed with a Marianas confocal microscope (Leica) with a 63x objective. For heat shock live imaging experiments 48 h post transfection, the coverslip was transferred to a FCS2 chamber assembled according to the manufacturer's instructions (Bioptechs). Media was perfused through the chamber, and then the chamber was placed into a Marianas spinning disk confocal system with a stage-top incubator and 63x objective with an objective heater (Bioptechs), both preheated to 37°C. The Microaqueduct Slide heater (FCS2 system) and the heated objective with 37°C immersion oil (Zeiss) were used to control the temperature. Movies were collected on a Marianas confocal microscope with 63x objective, with 40-sec intervals for the live imaging experiments, 200-ms intervals for the FLAP experiments of the time period. Average and standard errors were calculated from 3 independent experiments measuring at least 35 cells.

ATP measurement

Cellular contents of ATP were measured using CellTiter-Glo 2.0 assay kit (Promega; G9242) according to the manufacturer's instructions. 200 mM 2-deoxy-D-gluose (Millipore Sigma; D6134) was used to inhibit glycolysis pathway.

Statistical analysis

Statistical analysis was performed in GraphPad Prism. Comparisons between two means were performed by two-tailed *t* test. Comparisons among multiple means over 2 were performed by one-way ANOVA with Tukey's test. Mantel-Cox test was used to compare the dissociation curves showing cells with stress granules in live-cell imaging.

**Table S1. (separate file) Integrated stress granule proteome and VCP adaptors.** Table showing a list of proteins identified in stress granules and integrated stress granule proteome used in Fig. 5A. It also contains a list of 35 VCP adaptors.

**Movie S1. Impaired disassembly of heat shock-induced stress granules by TAK-243.** U2OS GFP-G3BP1 cells in the presence or absence of TAK-243 were incubated at 37°C for 2 min, 43°C for 30 min, and 37°C for 88 min, and GFP signals were monitored with 30-sec intervals. See Fig. 1, M and N.

**Movie S2. Impaired disassembly of heat shock-induced stress granules composed of G3BP1 6KR or 10KR.** U2OS *G3BP1/2* dKO cells stably expressing GFP-G3BP1 WT, NTF2L 6KR, RRM 4KR, or NTF2L/RRM 10KR were incubated at 37°C for 2 min, 43°C for 60 min, and 37°C for 118 min, and GFP signals were monitored with 30-sec intervals. See Fig. 2, G and H.

**Movie S3. Impaired disassembly of heat shock-induced stress granules by loss of ATP.** U2OS GFP-G3BP1 cells were incubated at 37°C for 2 min, 43°C for 60 min, and 37°C for 118 min, and GFP signals were monitored with 30-sec intervals. See fig. S3, D and E.

**Movie S4. Impaired disassembly of heat shock-induced stress granules by CB-5083.** U2OS GFP-G3BP1 cells in the presence or absence of CB-5083 were incubated at 37°C for 2 min, 43°C for 60 min, and 37°C for 118 min, and GFP signals were monitored with 30-sec intervals. See Fig. 4, A and B.

**Movie S5. Impaired disassembly of heat shock-induced stress granules by gene silencing of VCP.** U2OS GFP-G3BP1 cells were transfected with non-targeting siRNA or VCP siRNA and incubated at 37°C for 2 min, 43°C for 60 min, and 37°C for 118 min, and GFP signals were monitored with 30-sec intervals. See Fig. 4, C and D.

**Movie S6. Impaired disassembly of heat shock-induced stress granules by ALS-associated VCP mutants.** U2OS GFP-G3BP1 cells were transfected with VCP-GFP WT, R155H, or A232E and incubated at 37°C for 2 min, 43°C for 60 min, and 37°C for 118 min, and GFP signals were monitored with 30-sec intervals. See Fig. 4, E and F.

**Movie S7 and S8. Limited redistribution of G3BP1 from stress granules by VCP knockdown.**
U2OS cells stably expressing photoactivatable G3BP1 (PAGFP-G3BP1) were transfected with non-targeting siRNA (S7) or VCP siRNA (S8). 48 h post transfection, ROIs were activated by 405-nm laser and GFP signals were monitored with 200-ms intervals. See Fig. 4, G and H.

**Movie S9. Impaired disassembly of heat shock-induced stress granules by FAF2 knockdown.** U2OS GFP-G3BP1 cells were transfected with non-targeting siRNA or FAF2 siRNA and incubated at 37°C for 2 min, 43°C for 60 min, and 37°C for 118 min, and GFP signals were monitored with 30-sec intervals. See Fig. 5, E and F.

#### References and Notes


1. P. Yang *et al.*, G3BP1 Is a Tunable Switch that Triggers Phase Separation to Assemble Stress Granules. *Cell* **181**, 325-345 e328 (2020).
2. E. Tresse *et al.*, VCP/p97 is essential for maturation of ubiquitin-containing autophagosomes and this function is impaired by mutations that cause IBMPFD. *Autophagy* **6**, 217-227 (2010).
3. K. Zhang *et al.*, Stress Granule Assembly Disrupts Nucleocytoplasmic Transport. *Cell* **173**, 958-971 e917 (2018).
4. M. D. Figley, G. Bieri, R. M. Kolaitis, J. P. Taylor, A. D. Gitler, Profilin 1 associates with stress granules and ALS-linked mutations alter stress granule dynamics. *J Neurosci* **34**, 8083-8097 (2014).

### Figure S1

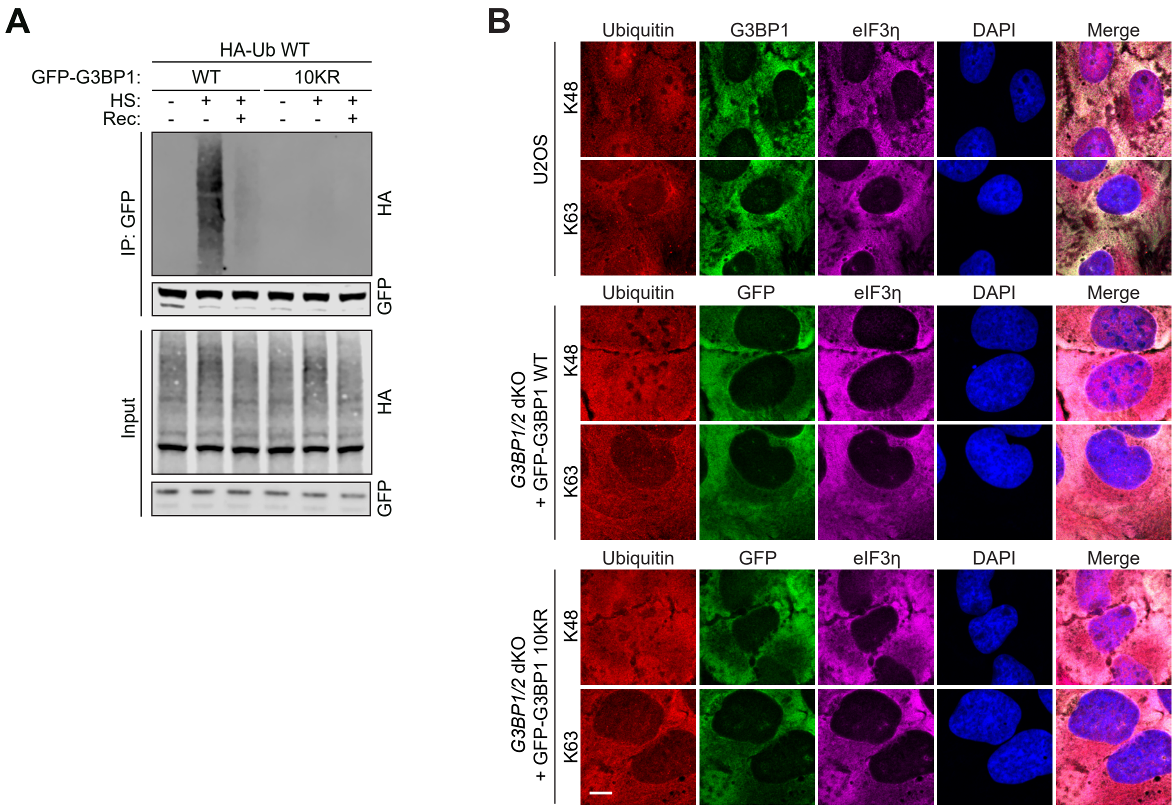


**Fig. S1. G3BP1 undergoes K63-linked ubiquitination in response to heat stress.** (**A**) Western blot of HEK293T cells transfected with HA-Ub and either GFP-G3BP1 WT or NTF2L/RRM KR (10KR) mutant constructs. Cells were exposed to no stress, heat shock (43°C, 1 h), or heat shock plus recovery (43°C, 1 h; 37°C, 30 min). Cell extracts were captured with magnetic beads conjugated with GFP antibody for IP and resulting beads were analyzed by immunoblot. (**B**) Control images corresponding to Figure 1I. Scale bar, 10 µm.

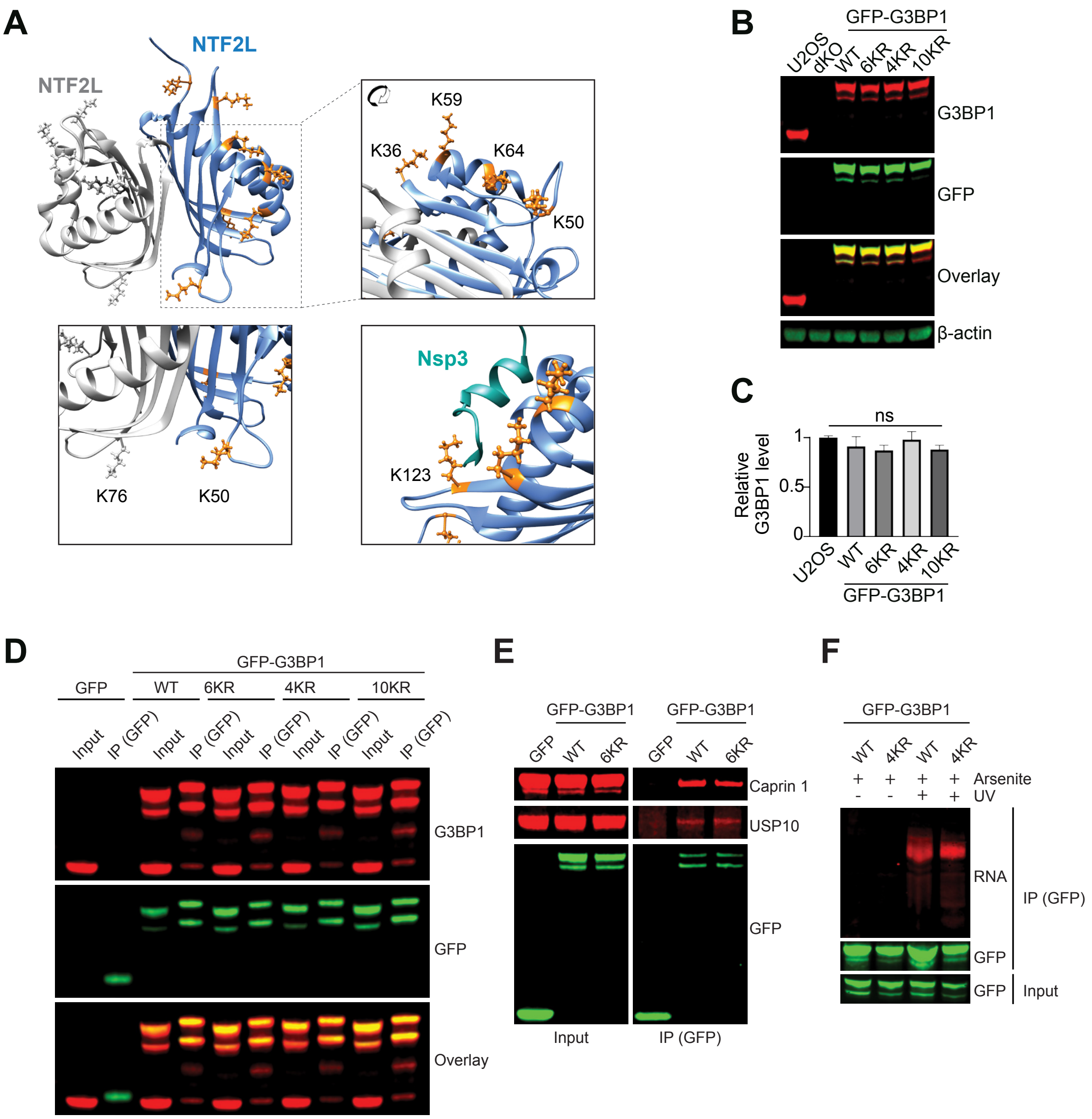


**Fig. S2. Generation of stable cell lines expressing G3BP1 mutant proteins; ubiquitination does not influence G3BP1 mobility during stress granule assembly.** (**A**) Dimeric structure of G3BP1 NTF2L domain enlarged with K36, K50, K59, and K64 cluster (upper right), K50 and K76 from each NTF2L domain (lower left), and K123 and hydrophobic grove where nsP3 protein of Old World alphaviruses makes a contact (lower right). (**B-C**) Western blot of U2OS G3BP1/2 dKO cells transfected with GFP-G3BP1 WT, 6KR, 4KR, or 10KR. Quantification of immunoblots from 3 replicates is shown in (**C**). Error bars indicate s.e.m. ns, not significant, ANOVA with Tukey's test. (**D**) Dimeric interaction of GFP-G3BP1 WT and KR mutants with endogenous G3BP1. HEK293T cells were transfected with pEGFP-C3 or indicated GFP-G3BP1 constructs. Cell extracts were captured with magnetic beads conjugated with GFP antibody for IP and resulting beads were analyzed by immunoblot. (**E**) Interaction of GFP-G3BP1 NTF2L 6KR mutant with Caprin1 and USP10. HEK293T cells were transfected with pEGFP-C3, GFP-G3BP1 WT or NTF2L 6KR constructs. Cell extracts were captured with magnetic beads conjugated with GFP antibody for IP and resulting beads were analyzed by immunoblot. (**F**) Crosslinking of RNA to both G3BP1 WT and RRM KR mutant. HEK293T cells were transfected with GFP-G3BP1 WT or RRM 4K mutant. Cells were exposed to no stress or oxidative stress (0.5 mM sodium arsenite, 1 h) and protein-RNA complexes were crosslinked with UV. Cell extracts were captured with magnetic beads conjugated with GFP antibody for IP and resulting beads were analyzed by immunoblot.

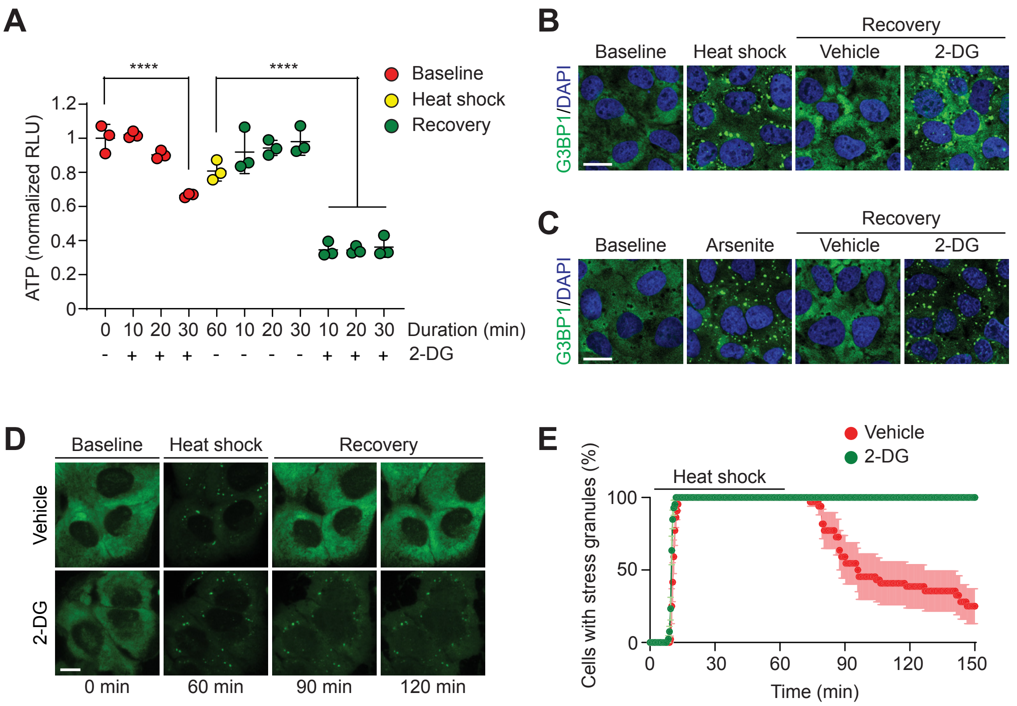


**Fig. S3. ATP is required for the disassembly of stress granules.** (**A**) ATP assay of U2OS cells were exposed to no stress, heat shock (43°C, 1 h) or indicated duration of 37°C recovery, in the presence or absence of 200 mM 2-deoxy-D-glucose (2-DG). Assay shows reduced cellular ATP contents by 2-DG during recovery after heat shock. Error bars indicate s.e.m. ****P< 0.0001, ANOVA with Tukey's test. (**B**) Fluorescent imaging of U2OS cells exposed to no stress, heat shock (43°C, 1 h), or heat shock plus recovery (43°C, 1 h; 37°C, 1 h) in the presence or absence of 200 mM 2-DG. Images show impaired disassembly of stress granules with decreasing cellular ATP. Scale bar, 20 µm. (**C**) Fluorescent imaging of U2OS cells exposed to no stress, oxidative stress (0.5 mM sodium arsenite, 1 h) or recovery with culture media (1 h) in the presence or absence of 200 mM 2-DG. Images show impaired disassembly of stress granules with decreasing cellular ATP. Scale bar, 20 µm. (**D-E**) Live-cell imaging of U2OS cells stably expressing GFP-G3BP1. Cells were incubated at 37°C for 2 min, 43°C for 60 min, and 37°C for 118 min, and GFP signals were monitored with 30-sec intervals. 200 mM 2-DG or the same volume of dissolving media were added to cells 62 min after monitoring, when the temperature returned to 37°C. Images show impaired disassembly of heat shock-induced stress granules. Representative still images are shown in (D). The percentage of cells with ≥ 2 stress granules is plotted in (E). Scale bar, 10 µm. Error bars indicate s.e.m. See also Movie S3.

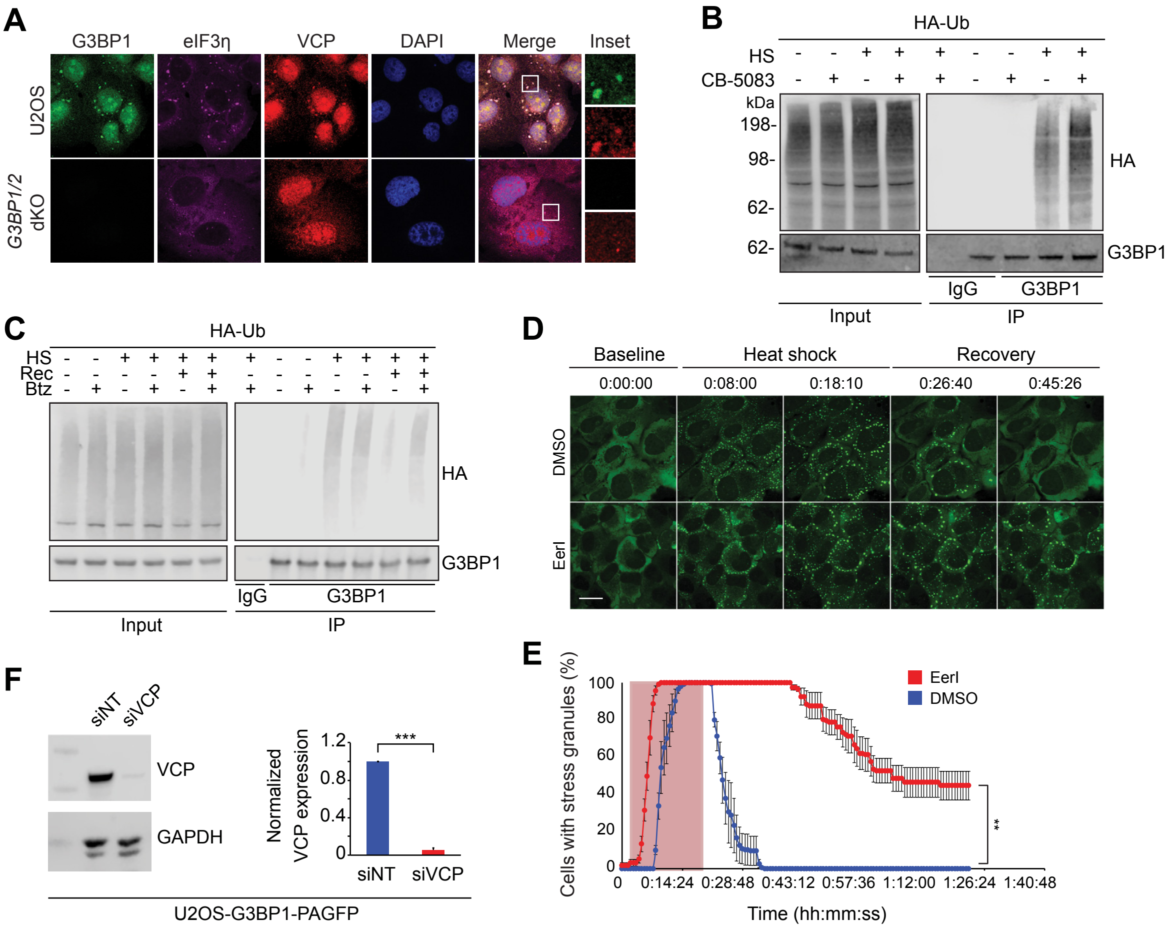


**Fig. S4. Impairing VCP function perturbs stress granule dynamics.** (**A**) Fluorescent imaging of U2OS and U2OS G3BP1/2 dKO cells exposed to heat shock (43°C, 1 h). Scale bar, 20 μm. (**B**) Western blot of HEK293T cells transfected with HA-tagged ubiquitin, treated with DMSO or CB-5083 (1 h) and exposed to heat shock (43°C, 1 h). Cell extracts were captured with magnetic beads conjugated with G3BP1 antibody for IP and resulting beads were analyzed by immunoblot. Blots suggest VCP-mediated decoupling of ubiquitin chains anchored to G3BP1 in response to heat shock. (**C**) Western blots of HEK293T cells transfected with HA-tagged ubiquitin and exposed to no stress, heat shock (43°C, 1 h), or heat shock plus recovery (43°C, 1 h; 37°C, 1 h) in the presence or absence of bortezomib (Btz). Cell extracts were captured with magnetic beads conjugated with G3BP1 antibody for IP and resulting beads were analyzed by immunoblot. Blots suggest that ubiquitinated G3BP1 is targeted to the proteasome upon recovery. (**D-E**) Live-cell imaging of U2OS cells stably expressing GFP-G3BP1 and treated with DMSO or eeyarestatin I (EerI, 56 μM) for 30 min prior to imaging. Cells were exposed to heat shock (43°C, 18 min), then allowed to recover for indicated times. GFP signals were monitored with 40-sec intervals. Scale bar, 10 μm. The percentage of cells with ≥ 2 stress granules is plotted in (E). Scale bar, 20 μm. Error bars indicate s.d. **P< 0.01, Mantel-Cox test. (**F**) Western blots of U2OS cells stably expressing PAGFP-G3BP1 and transfected with either non-targeting siRNA (siNT) or VCP siRNA. Cell were harvested 48 h post transfection. ***P< 0.001, student t-test.

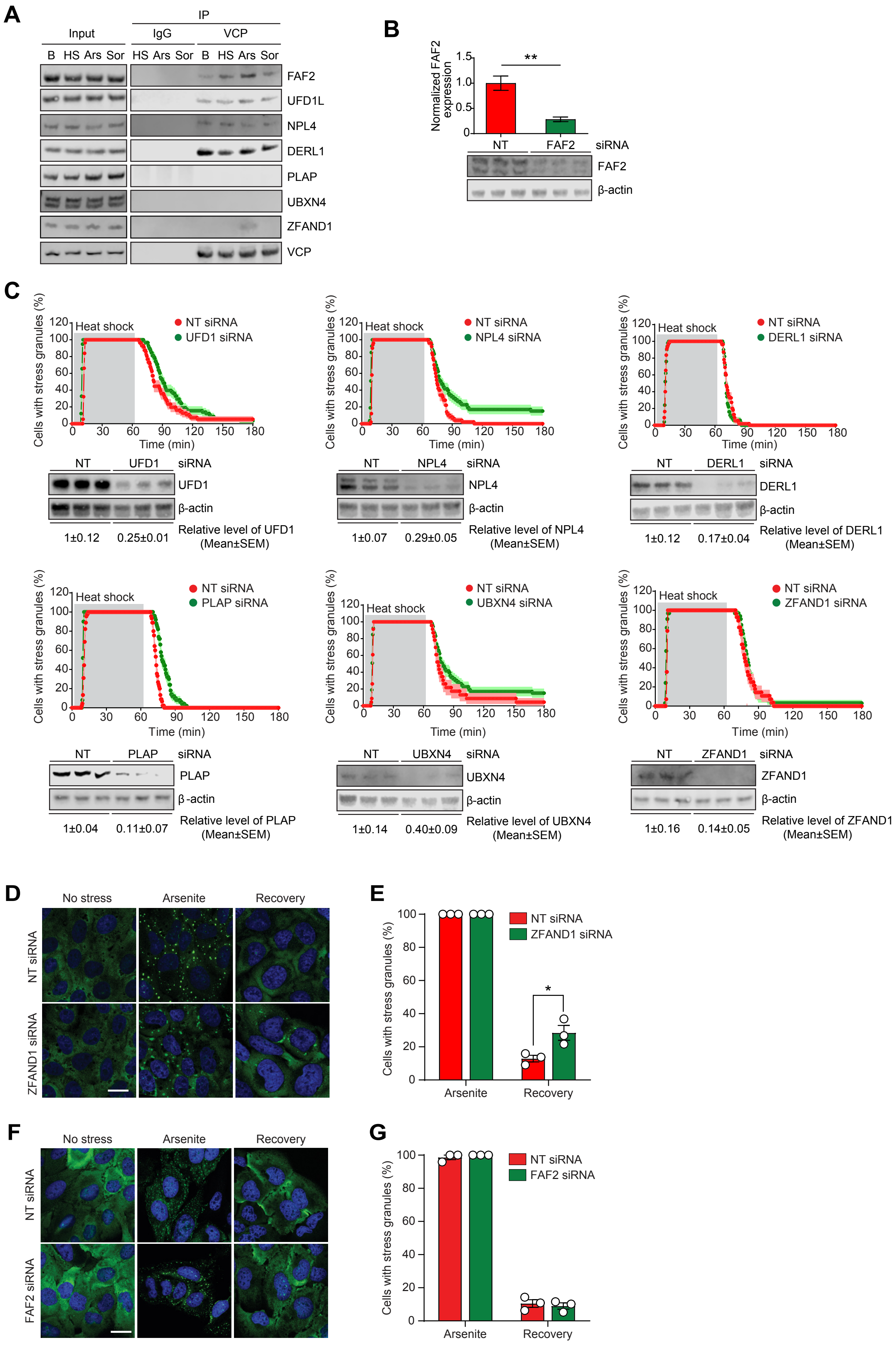

A
IP
Input
IgG
VCP
B HS Ars Sor HS Ars Sor B HS Ars Sor
FAF2
UFD1L
NPL4
DERL1
PLAP
UBXN4
ZFAND1
VCP
B
Normalized FAF2 expression
**
NT
FAF2
siRNA
FAF2
β-actin
C
Cells with stress granules (%)
Heat shock
NT siRNA
UFD1 siRNA
Time (min)
NT
UFD1
siRNA
UFD1
β-actin
1±0.12
0.25±0.01
Relative level of UFD1 (Mean±SEM)
NT siRNA
NPL4 siRNA
NPL4
β-actin
1±0.07
0.29±0.05
Relative level of NPL4 (Mean±SEM)
NT siRNA
DERL1 siRNA
DERL1
β-actin
1±0.12
0.17±0.04
Relative level of DERL1 (Mean±SEM)
NT siRNA
PLAP siRNA
PLAP
β-actin
1±0.04
0.11±0.07
Relative level of PLAP (Mean±SEM)
NT siRNA
UBXN4 siRNA
UBXN4
β-actin
1±0.14
0.40±0.09
Relative level of UBXN4 (Mean±SEM)
NT siRNA
ZFAND1 siRNA
ZFAND1
β-actin
1±0.16
0.14±0.05
Relative level of ZFAND1 (Mean±SEM)
D
No stress
Arsenite
Recovery
NT siRNA
ZFAND1 siRNA
E
Cells with stress granules (%)
NT siRNA
ZFAND1 siRNA
*
Arsenite
Recovery
F
No stress
Arsenite
Recovery
NT siRNA
FAF2 siRNA
G
Cells with stress granules (%)
NT siRNA
FAF2 siRNA
Arsenite
Recovery

**Fig. S5. FAF2 links ubiquitinated G3BP1 to VCP.** (**A**) Western blot of U2OS cells exposed to no stress, heat shock (43°C, 1.5 h), oxidative stress (0.5 mM sodium arsenite, 1.5 h), or osmotic stress (0.4 M sorbitol, 1.5 h). Cell extracts were captured with magnetic beads conjugated with VCP antibody for IP and resulting beads were analyzed by immunoblot. Blots assess interaction between VCP and stress granule-resident adaptors described in Figure 5A. (**B**) Western blot of U2OS cells transfected with non-targeting (NT) siRNA or FAF2 siRNA for 48 h. Blots show knockdown efficiency of FAF2 siRNA as used in Figure 5. **$P< 0.01$, two-tailed t test. (**C**) Graphs show results from fluorescent imaging of U2OS cells stably expressing GFP-G3BP1 and transfected with non-targeting siRNA or siRNAs of indicated genes (all stress granule-resident VCP adaptors). Cells were incubated at 37°C for 2 min, 43°C for 60 min, and 37°C for 118 min, and GFP signals were monitored with 30-sec intervals. The percentage of cells with ≥ 2 stress granules is plotted. Western blots below each graph demonstrate knockdown of indicated genes; each condition is shown in triplicate. Blots were measured with densitometry and relative expression levels are shown below each blot. (**D-E**) Fluorescent imaging of U2OS cells exposed to no stress, oxidative stress (0.5 mM sodium arsenite, 1 h) or recovery with culture media (1 h) after transfection of non-target (NT) siRNA or ZFAND1 siRNA. Scale bar, 20 µm. Quantification of cells with stress granules from 3 replicates is shown in (E). Error bars indicate s.d. *$P< 0.05$, ANOVA with Tukey's test. (**F-G**) Fluorescent imaging of U2OS cells exposed to no stress, oxidative stress (0.5 mM sodium arsenite, 1 h) or recovery with culture media (1 h) after transfection of non-target (NT) siRNA or FAF2 siRNA. Scale bar, 20 µm. Quantification of cells with stress granules from 3 replicates is shown in (G). Error bars indicate s.d. ns, not significant, ANOVA with Tukey's test.

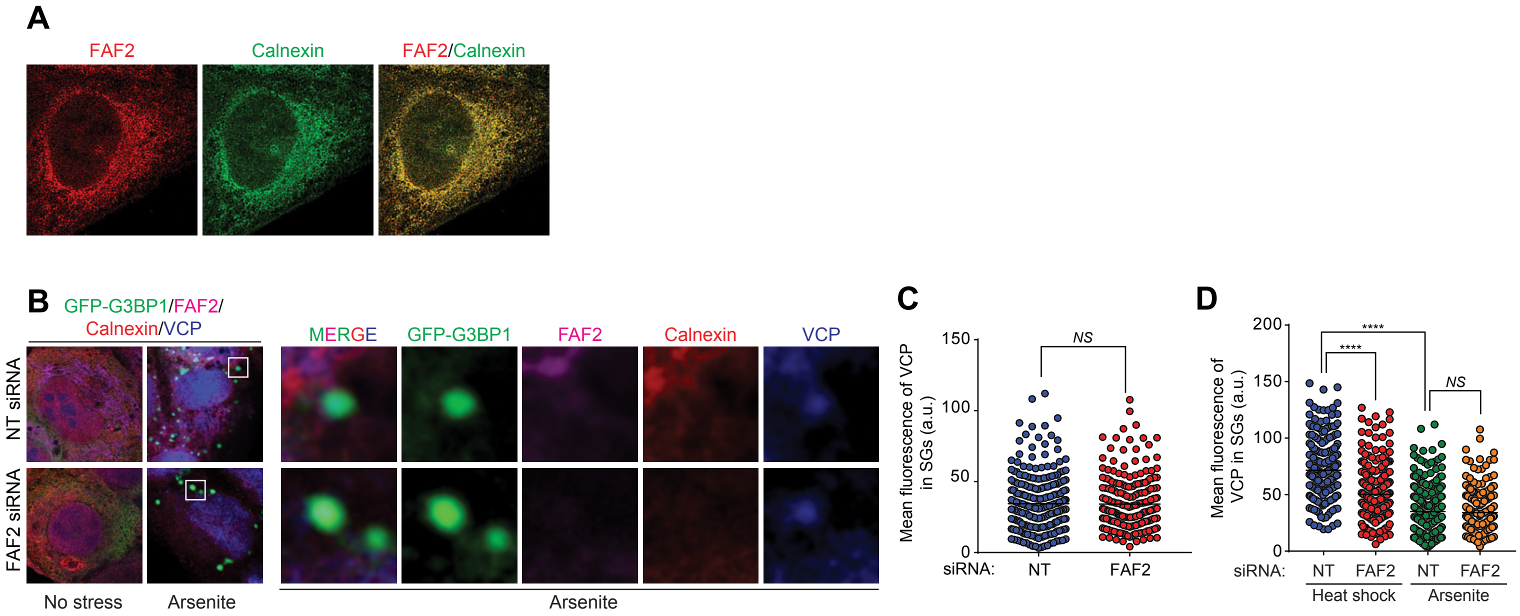


**Fig. S6. FAF2, an ER-resident VCP adaptor, links ER to G3BP1 for disassembly of stress granules upon heat shock.** (**A**) ER localization of FAF2. Unstressed U2OS cells were fixed with FAF2 and calnexin antibodies for IF. (**B-C**) Imaging of U2OS cells stably expressing GFP-G3BP1 and exposed to no stress or 0.5 mM sodium arsenite, fixed, and stained with indicated antibodies for IF. Fluorescent intensities of VCP in stress granules are plotted in (C). Result show that recruitment of VCP to arsenite-induced stress granules is not regulated by FAF2. Error bars indicate s.e.m. ns, not significant, two-tailed t test. (**D**) Combined plots of Figure 5H and Figure S5F for comparison of VCP recruitment to stress granules. Error bars indicate s.e.m. ns, not significant, ****P< 0.0001, ANOVA with Tukey's test.
